# Improving the preclinical and clinical success rates of LMW drugs depends on radical revisions to the status quo scientific foundations of medicinal chemistry: a case study on COVID M^pro^ inhibition

**DOI:** 10.1101/2022.10.31.514572

**Authors:** Robert A. Pearlstein, Hongbin Wan, Sarah Williams

**Author notes:** Corresponding author: Robert A. Pearlstein, Ph.D. Phone: -871-7293. These authors contributed equally to this work (Robert A. Pearlstein, Hongbin Wan).

## Abstract

The poor preclinical and clinical success rates of low molecular weight (LMW) compounds can be partially attributed to the inherent trial-and-error nature of pharmaceutical research, which is limited largely to <u>retrospective</u> data-driven, rather than <u>prospective</u> prediction-driven human relevant workflows stemming from: 1) inadequate scientific understanding of structure-activity, structure-property, and structure-free energy relationships; 2) disconnects between empirical models derived from in vitro equilibrium data (e.g., Hill and Michaelis-Menten models) vis-à-vis the native non-equilibrium cellular setting (where the pertinent metrics consist of rates, rather than equilibrium state distributions); and 3) inadequate understanding of the non-linear dynamic (NLD) basis of cellular function and disease. We argue that the limit of understanding of cellular function/dysfunction and pharmacology based on <u>empirical</u> principles (observation/inference) has been reached, and that further progress depends on understanding these phenomena at the <u>first principles</u> theoretical level. Toward that end, we have been developing and applying a theory (called “Biodynamics”) on the general mechanisms by which: 1) cellular functions are conveyed by dynamic multi-molecular/-ionic (multi-flux) systems operating in the NLD regime; 2) cellular dysfunction results from molecular dysfunction; 3) molecular structure and function are powered by covalent/non-covalent forms of free energy; and 4) cellular dysfunction is corrected pharmacologically. Biodynamics represents a radical departure from the status quo empirical science and reduction to practice thereof, replacing: 1) the interatomic contact model of structure-free energy and structure-property relationships with a solvation free energy model; 2) equilibrium drug-target occupancy models with dynamic models accounting for time-dependent drug and target/off-target binding site buildup and decay; and 3) linear models of molecular structure-function and multi-molecular/-ionic systems conveying cellular function and dysfunction with NLD models that more realistically capture the emergent non-linear behaviors of such systems. Here, we apply our theory to COVID M^pro^ inhibition and overview its implications for a holistic, in vivo relevant approach to drug design.

## Introduction

Drug discovery is analogous to navigating a multi-dimensional maze with multiple entrances (i.e., targets + chemical starting points), few exits (i.e., approved drugs), and many blind alleys in between. Successful navigation depends on finding one or more paths to a safe therapeutic index (the ratio of efficacious to toxic drug exposure) in humans, the overall odds of which (P_d_) are determined by the following sets of conditional probabilities:

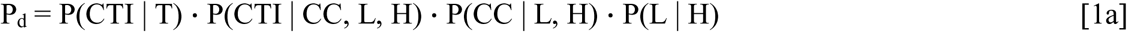

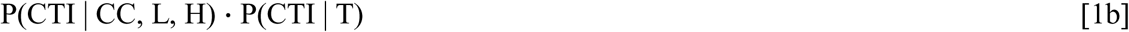

where T is a biomolecular target on the path to efficacy, CTI is a clinical candidate for which a safe therapeutic index (TI) is achieved in clinical testing, CC are clinical candidates on the path to the CTI, L is a lead series on the path to the CC, H are hits on the path to L, and P(x_n_ | x_n+1_, …, x_n+m_) are the conditional probabilities of success at each step of the process (e.g., CC depends on L, which depends on H). P_d_ is low by all accounts (Wong et al., 2019) (although indeterminate on a case-by-case basis), which is partially attributable to heavy reliance on trial-and-error synthesis and testing workflows guided by poorly predictive experimental and computational models for target and off-target potency (structure-activity/structure-free energy), solubility, permeability, and pharmacokinetic (PK)/ADMET (absorption, distribution, metabolism, elimination, and toxicity) properties <u>relevant to the human</u> <u>setting</u>. Scaling the number of successful outcomes at the current P_d_ depends on scaling the number of attempts, whereas improving the underlying P(x_n_ | x_n+1_, …, x_n+m_) depends on improving the predictiveness of the aforementioned models.

All scientific disciplines are broadly comprised of empirical and theoretical/first principles branches. Empirical science is based on observations of “what”, “where”, and “when” (the parts, systems, and behaviors thereof), the “how” and “why” (the mechanisms), which are inferred anecdotally or from statistical correlations with other observations that lead in some cases to empirical “laws”. Theoretical science is based on deductive/logical reasoning that extends beyond observation, whereas empirical science depends heavily on technologies that extend the powers of observation, curation, and data analysis. The major scientific theories, including special/general relativity, quantum mechanics, the standard model of particle physics, and number theory, originated from physics and math. Conversely, chemistry (with the exception of quantum chemistry) is a largely empirical science that revolves around data models that are incorrect, inadequate, or right for the wrong reasons when:

1. Fit to noisy or under-sampled data (noting that weak signals can be extracted from low signal-to-noise data using machine learning or artificial intelligence when actually present in the data).
2. Fit using descriptors/parameters that poorly explain the data, often resulting in overfitting (i.e., a disproportionate number of descriptors/parameters relative to the size of the dataset).
3. Fit to data measured in one context and extrapolated to another.
4. Predicated on scalar rather than vector quantities needed to explain multi-dimensional phenomena.

The use of empirical models in contexts other than those in which they were derived can result in incorrect predictions and inferences for the aforementioned reasons (e.g., Newton’s empirical gravitational law predicts well except under conditions of extreme gravity, whereas Einstein’s general relativity theory predicts well except in black holes). In this work, we:

1. Revisit the “how” and “why” of cellular function conveyed by multi-molecular and multi-ionic systems, cellular dysfunction caused by dysfunctional biomolecules, and pharmacodynamics (PD) through the lens of a novel first principles multi-scale theory called “Biodynamics” that we have been developing, testing, and using to study a wide range of phenomena over the last several years (Pearlstein et al., 2017, 2021; Selvaggio and Pearlstein, 2018; Selvaggio et al., 2020; Wan et al., 2020a), including the structure-function of cereblon (Wan et al., 2021a), G-protein coupled receptors (GPCRs), voltage-gated ion channels (Wan and Pearlstein, 2022), and COVID M^pro^ (Wan et al., 2020a), as well as hERG blockade (Wan et al., 2020b, 2021b) and cellular arrhythmia (Selvaggio et al., 2020).
2. Offer an integrated top-down perspective on molecular structure-function/dysfunction, drug-target/off-target binding, PK, PD, and non-covalent free energy within and between the atomic (zoomed-in) and systems (zoomed-out) levels and examine the implications of this for improving P_d_ based on in vivo relevant drug design.
3. Revisit our previous work on COVID M^pro^, focusing on the structure-binding free energy relationships of the inhibitors nirmatrelvir and an analog thereof (PF-00835231).

### Toward a first principles understanding of the “how” and “why” of living cells

The moving parts of living cells consist entirely of molecules and ions organized into multi-molecular/ionic systems exhibiting non-random, <u>non-linear time-dependent</u> behaviors (commonly referred to as “complex” systems) (Noble, 2006). In fact, nearly all natural systems comprised of three or more moving parts, ranging from subatomic particles to atoms, molecules, the atmosphere, oceans, stars, the solar system, and the Universe itself behave non-linearly (whereas linear systems may be comprised of thousands of moving parts). All such systems are powered by attractive and repulsive forms of energy that are counter-balanced within a Goldilocks zone in which both the bound state (global stability) and rearrangeability (local instability) are maintained over time (e.g., gravity vis-à-vis fusion in the case of stars, gravity vis-à-vis kinetic energy in the case of the solar system, and electromagnetic attraction vis-à-vis kinetic energy in the case of atoms and covalent bonds). Science in general revolves around observing, describing, explaining, and predicting the configurational states and behaviors of natural NLD systems (many of which are strange and seemingly magical). Such systems:

1. **“Synthesize/solve” their overall behaviors (commonly referred to as emergence)**, which in all cases, exceed the sum of the behaviors of their constituent parts (where the behavior of each part depends on the behavior of the entire system, and the behavior of the entire system depends on the behaviors of the parts and their mutual interactions). As such, NLD behaviors can only be observed or simulated, but not predicted via analytical math (i.e., they are deterministically unpredictable).
2. **Exhibit chaotic behaviors at perturbation-induced tipping points** (e.g., cardiac arrhythmias caused by perturbation of the normal dynamic inward-outward ion current balance; ice ages caused by perturbation of the normal dynamic atmospheric-oceanic-geo-thermal balance; implosions/explosions of stars caused by the loss of dynamic balance between gravity and fusion-driven thermal expansion).
3. **Respond to perturbations in a non-causal fashion**, where causes are effects and effects are causes.
4. **Are powered by non-equilibrium flows between high and low energy states** (referred to as “sources” and “sinks”, respectively), analogous to the flow of air from high to low pressure regions of the atmosphere. The molecular populations of cellular systems build and decay via synthesis (sources) and degradation (sinks), as do their non-covalently bound states, which are powered within a Goldilocks zone of favorable and unfavorable enthalpic and entropic energy contributions.

Living cells comprise a unique class of natural NLD systems in that their moving parts consist exclusively of micro, nano, pico, or femto quantities of diverse ions and LMW and high molecular weight (HMW) species operating on the microscopic length and time scales (subserving high density information storage and fast information processing), versus bulk quantities of molecules operating on the geologic length and time scales (e.g., the atmosphere), and heavenly bodies operating on the cosmic length and time scales. Cells “solve” their behaviors <u>at each instant of time</u> via a physical form of math commonly known as “analog computing”, in which the hardware (molecules and ions) and software (NLD operation of the system) are one and the same. The normal and abnormal behaviors of macromolecular constituents and multi-molecular/multi-ionic systems conveying cellular functions cannot be understood or reliably predicted via empirical models based on equilibrium data measured for the isolated parts (equating to tacit linearization of such systems) or even intact (typically immortal) cultured cell lines. Further increases in P(CTI | T) and P(CTI | CC, L, H) depend on connecting the dots between target selection and clinical candidate generation (the in vitro ➔ native cellular scenarios).

Improved predictions of biological behaviors and the effects of drugs thereon depend on a first principles theoretical understanding of the cellular “playbook” shared among all natural NLD systems, including the means by which they are powered. The reduction to practice of Biodynamics consists of first principles analyses and simulations of:

1. Solvation free energy, which we postulate is the principal form of free energy powering the <u>non-covalent</u> inter- and intramolecular rearrangements of aqueous solutes (Wan et al., 2020a), versus interatomic contact free energy under the status quo paradigm (noting that improving P(CC | L, H) and P(L | H) depends heavily on a correct understanding of structure-free energy relationships). Under non-equilibrium cellular conditions, intermolecular non-covalent free energy consists of association and dissociation barriers (denoted as 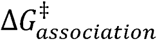 and 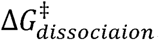) to which the rate constants k_on_ and k_off_ are proportional, and intramolecular non-covalent free energy consists of entry and exit barriers (denoted as 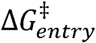 and 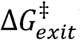) to which the rate constants k_in_ and k_out_ are proportional. We proposed in 2010 that 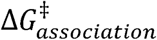 and 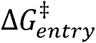 consist principally of the total desolvation cost incurred in entering new inter- and intramolecular states and 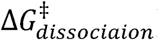 and 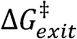 consist principally of the resolvation cost incurred in exiting those states (Pearlstein et al., 2010, 2013; Tran et al., 2013, 2014; Velez-Vega et al., 2014; Wan et al., 2020a, 2020b, 2021b, 2021a; Wan and Pearlstein, 2022), an idea inspired by the pioneering work of McKay (Pearlstein et al., 2017) and Kurtzman (Young et al., 2007a; Abel et al., 2008) on the microscopic behaviors of solvating water. Our work has led to novel insights about the general mechanisms by which free energy is transduced into molecular structure and function under non-equilibrium conditions, including the folding of HMW species and non-covalent binding between HMW-LMW and HMW-HMW species. Solvation free energy is qualitatively mirrored by the position-dependent occupancy of solvating water on the external surfaces of LMW solutes and the external and internal/buried surfaces of HMW solutes (noting that <u>scalar</u> solvation free energy contributions cannot be accurately predicted via force-field-based methods, or practically via quantum chemical methods). The <u>spatial distribution</u> of solvation free energy across a given solute surface relative to bulk solvent depends on the local H-bonding environment of each solvating water, the collective properties of which we refer to as the “solvation field” (noting that H-bond enriched, bulk-like, and H-bond depleted solvation are <u>statistically</u> distributed in a Gaussian fashion (Wan et al., 2020a)). The free energy barriers governing the rates of entry and exit to/from non-covalent structural states are determined by the degree of complementarity between the solvation fields residing at the binding interfaces of cognate partners, including drugs and their respective targets and off-targets (i.e., H-bond enriched matched to H-bond enriched and H-bond depleted matched to H-bond depleted). We refer to the fields in these regions as “solvophores” and “inverse solvophores”, borrowing from the pharmacophore-inverse pharmacophore concept, and the collection of solvophores and inverse solvophores across the entire endogenous space as the “solvome” (Wan et al., 2021a). Contrary to popular belief, solvation free energy is comprised of both enthalpic and entropic contributions based on the following rationale (Pearlstein et al., 2017):

a. The H-bond <u>enthalpy</u> of liquid water normalized to its effective 30 Å^3^ molecular volume is by far the highest among all liquid solvents and dissolved aqueous substances to the extent that the non-covalent solute structure and the dynamic properties thereof are powered principally by the minimization of water H-bond enthalpy at varying degrees of entropic cost (Pearlstein et al., 2017). Entropic losses relative to bulk solvent (taken as the zero-entropy reference) occur in <u>solvating</u> water in proportion to the degree of ordering required to achieve the minimum water-water and water-solute H-bond enthalpy (commonly referred to as entropy-enthalpy compensation).
2. The normal and abnormal behaviors of mutant and WT proteins in the context of the native NLD multi-molecular systems to which they belong, versus in isolation under in vitro conditions and in computational studies (Selvaggio et al., 2020; Pearlstein et al., 2021). The resulting models can, in principle, be used to <u>qualitatively</u> predict efficacious drug-target combinations with improved P(CTI | T). NLD behaviors at both the atomic (e.g., multi-body intra-protein and protein-water interactions during protein folding and binding), and multi-molecular system levels that convey cellular functions are implicit to our theory.
3. Non-covalent binding under non-equilibrium conditions, in which binding site levels/concentrations build and decay over time (which we refer to as “binding dynamics”) versus the status quo static equilibrium paradigm. As such, cellular functions are governed by rates (i.e., the fastest fluxes), rather than the equilibrium distributions of the participating species.

Lead optimization under the status quo paradigm is aimed at minimizing drug-target ΔG (reflected in equilibrium binding constants, including K_d_ and IC_50_), together with permeability, solubility, and other pharmacokinetic properties, in contrast to the Biodynamics approach, which is aimed at achieving a Goldilocks zone of balanced drug solvation free energy contributions governing target and off-target binding (corresponding to minimal drug and target <u>desolvation</u> costs underlying 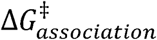 and maximal drug and target resolvation costs underlying 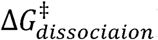), sufficient membrane permeability (corresponding to minimal drug and membrane desolvation costs and minimal drug and membrane resolvation costs needed to circumvent membrane partitioning), and sufficient solubility (which is directly proportional to solvation free energy) based on the following rationale:

1. ΔG applies only under equilibrium conditions, whereas 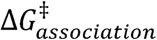 and 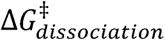 apply under both equilibrium and non-equilibrium conditions.
2. Structure is <u>physically</u> linked to the separate 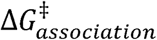 and 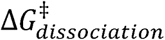 contributions (and 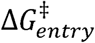 and 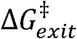 in the case of intramolecular rearrangements), rather than ΔG per se, which is merely the difference between these quantities (just as taxes are linked to income and expenses, rather than bank balances).
3. We showed that the minimum efficacious drug-target occupancy <u>at the lowest possible</u> <u>free C_max_</u> (denoted as γ_eff_) is achieved with <u>dynamic</u> binding sites when k_on_ is tuned to the rate of binding site buildup, and k_off_ is tuned to the rate of binding site decay (Selvaggio and Pearlstein, 2018), thereby reducing the risk of off-target binding, tox, adverse effects, and side effects that diminish P(CTI | CC, L, H).

Binding is thus governed by dynamic contributions analogous to a continual dance between the partners and solvating water, in which:

1. Association is accompanied by exchanges between the partners, their H-bond enriched, bulk-like, and H-bond depleted solvating water, and bulk solvent.
2. On-rate is given by k_on ⋅_ [free partner i](t) _⋅_ [free partner j](t), where k_on_ is proportional to the total cost of desolvating H-bond enriched water solvating the binding interface, which, in turn, is proportional to the quality of water-protein and water-water H-bond replacements by intermolecular H-bonds.
3. Dissociation is accompanied by exchanges between bulk solvent and the complex, resulting in the resolvation of H-bond enriched, bulk-like, and depleted positions of the dissociated partners.
4. Off-rate is given by k_off ⋅_ [partner i-partner j](t), where k_off_ is proportional to the total resolvation cost, which in turn, is proportional to the loss in number and/or strength of water H-bonds between bulk solvent (3.5 moderate H-bonds per water) and the solvation layer (0-4 weak H-bonds per water) (Pearlstein et al., 2017).
5. Water exchanges to/from bulk solvent and solvation incur:

a. Entropic gains and losses resulting respectively from water transfer to/from solvation and bulk solvent.
b. Enthalpic losses due to desolvation of H-bond enriched positions (which are partially offset by drug-target H-bond enthalpic gains) and gains due to the resolvation of H-bond enriched positions (which are partially offset by drug-target H-bond enthalpic losses).
c. Enthalpic gains due to desolvation of H-bond depleted positions and losses from resolvation of H-bond depleted positions.

### The implications of non-equilibrium NLD cellular behavior for medicinal chemistry, computational chemistry, and data science

We proposed in our previous work that cellular function is predicated on the notion of “physical math” (commonly known as analog computing), the “physical equations” of which consist of temporospatial changes in the concentrations or number densities of molecular species and the covalent and non-covalent intra- and/or intermolecular states thereof (which we refer to as “fluxes”) (Pearlstein et al., 2021; Wan et al., 2021a). Cellular functions (equating to emergent behaviors) are conveyed by specific sets of fluxes and static species that are coupled into systems via <u>transient</u> non-covalent binding interactions (Figure 1A). As mentioned above, the NLD behavior of such systems is holistic in that the behaviors of each flux (including drug PK and drug-target PD) depend on the behaviors of all other fluxes (i.e., the system) and vice versa, and as such, cannot be properly studied in isolation. Fluxes build and decay <u>exponentially</u>, and are therefore subject to runaway behavior (i.e., over- and under-shooting) unless sufficiently damped. We postulate that such behavior is circumvented by dynamic counterbalancing between flux-anti-flux pairs (“Yins” and “Yangs”) (Pearlstein et al., 2017, 2021), in which Yang buildups are “phased” relative to their corresponding Yins by “clocks” (with the intervening steps in the pathway equating to “ticks”) (Figure 1B). We further postulate that many cellular diseases result from over- or undershooting the functional levels of fluxes or their covalent or non-covalent states due to mutation-induced Yin-Yang imbalances caused by abnormal changes in the rates of Yin or Yang buildup or decay.

**Figure 1.**
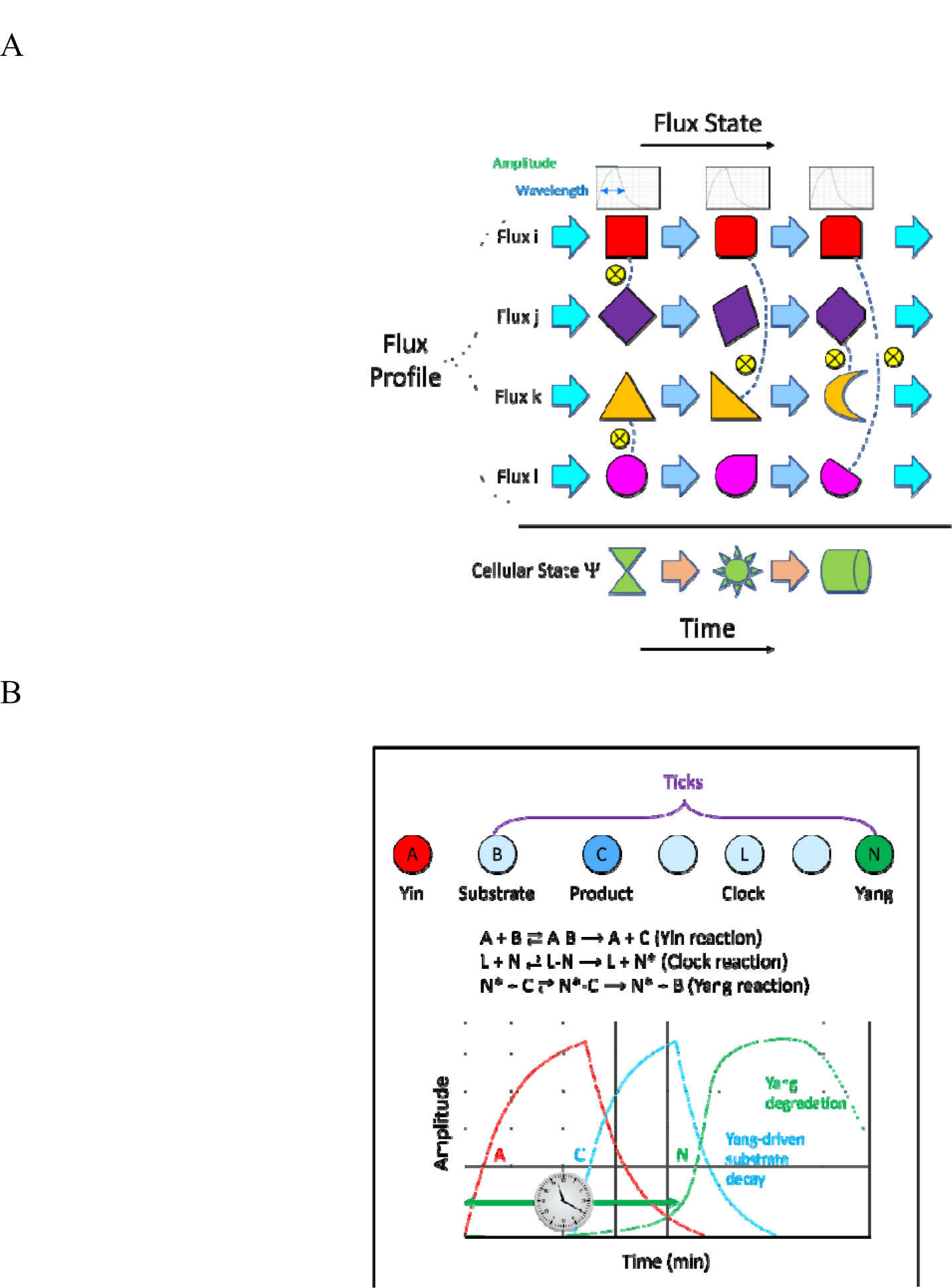
(A) Cellular functions are conveyed by systems comprised of multiple coupled molecular or ionic (in the case of action potentials) fluxes operating in the NLD regime, where each flux undergoes one or more non-covalent intra- and/or intermolecular and often one or more covalent state transitions (e.g., phosphorylation, ubiquitylation). Fluxes build and decay exponentially in time via synthesis and degradation or covalent activation and deactivation. (B) Runaway exponential behavior is damped via dynamic counterbalancing between flux (“Yin”)- anti-flux (“Yang”) pairs. Yin-Yang phasing is governed by “clocks” that drive Yang buildups downstream of their respective Yins, with the intervening steps equating to “ticks” (Pearlstein et al., 2021).

Non-covalent binding between fluxes i and j or flux i and static species j and enzymatic turnover are physical processes (“molecular differential equations” (MDEs)), the symbolic forms of which consist of the following:

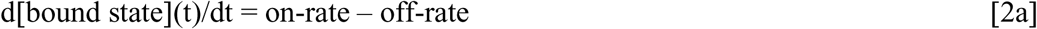

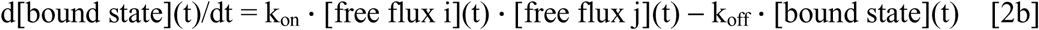

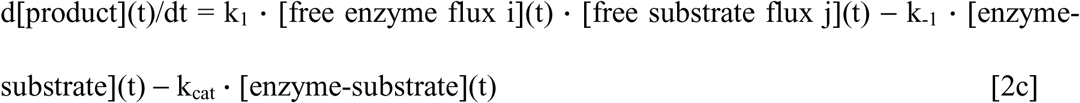

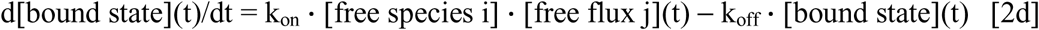

where d[bound state](t)/dt denotes the difference between the on- and off-rates (equation 2a), equating to the rates of buildup and decay of the bound state of a cognate flux pair (equations 2b-2c) or a flux and its static cognate partner (equation 2d). d[bound state](t)/dt builds when on-rate > off-rate, decays when off-rate > on-rate, and remains constant when on-rate = off-rate (noting that the net decay rate is determined, in part, by re-binding, as reflected in the on-rate).

The special case of drug-target binding is given symbolically by:

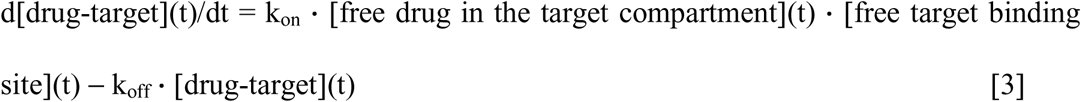

where d[free target binding site](t)/dt may vary from 0 (constant levels) to >> 0, case-by-case, and k_on_ determines the amount of free drug that can be “absorbed” by the target per unit time (where the excess drug “overflows” the binding site) (Selvaggio and Pearlstein, 2018; Pearlstein et al., 2021). Equations 2d and 3 contrast with the equilibrium free concentration-occupancy relationship given by the Hill equation:

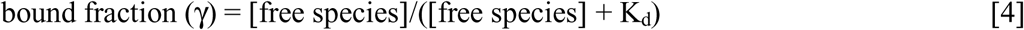

In our previous work, we showed that (Selvaggio and Pearlstein, 2018):

1. The solution of equation 3 converges to the steady state level (described by equation 4) when parity exists between k_on_ and k_off_ and the rates of binding site buildup (denoted k_i_) and decay (denoted k_-i_), respectively, which we refer to as “kinetically tuned binding” (Figure 2A).
2. The solution of equation 3 is less than the steady state level, such that drug-target occupancy (γ(t)) lags the dynamic free target concentration at all time points when k_on_ and k_off_ are mistuned to k_i_ and k_-i_, respectively (in proportion to the degree of mistuning) (Figure 2B).

**Figure 2.**
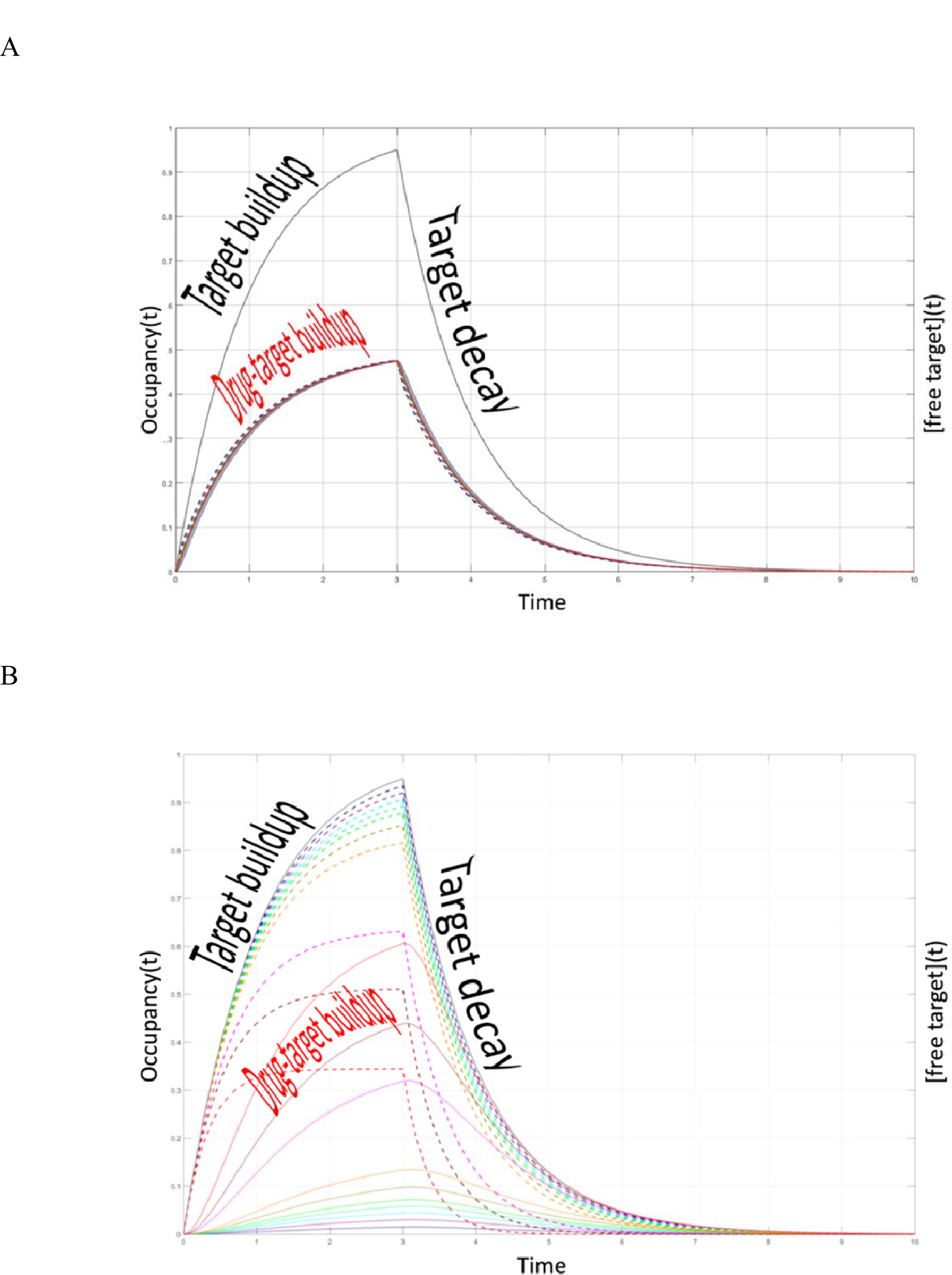
Hypothetical <u>simulated</u> target and drug-target γ(t) buildup/decay curves for a given k_i_, k_-i_, k_on_, and k_off_ (Selvaggio and Pearlstein, 2018). The bound and free drug concentrations are represented by the solid and dotted tracings, respectively. (A) The kinetically tuned scenario in which parity exists between k_on_ and k_i_ and k_off_ and k_-i_. Steady state γ(t) = 50% of [free target](t) is achieved at all time points when [free drug] = K_d_, and any given occupancy is achieved in multiples of K_d_. (B) The kinetically mistuned scenario in which parity between k_on_ and k_i_ and/or k_off_ and k_-i_ is absent. In this scenario, non-steady state γ(t) < steady state γ(t) occurs at each time point, such that [drug-target](t) buildup lags [free target](t) buildup. The expected minimum efficacious occupancy (denoted as γ_eff_) in humans is underestimated in such cases, the true level of which is only achievable during clinical testing via dose escalation. In the worst-case scenario, binding is <u>mistuned to the therapeutic target</u> and <u>tuned to one or more off-targets</u> by happenstance.

*Caveat 1: Kinetic mistuning results in disconnects between the true versus expected drug-targetγ(t) and minimum efficacious exposure in vivo, requiring dose escalation in the clinic to identify the true efficacious exposure (thereby eroding the safety margin, TI, and P(CTI | CC, L, H))*.

Pharmacological intervention is putatively aimed at restoring normal Yin-Yang counterbalancing (the PD response) of afflicted pathways via direct inhibition or activation of the dysfunctional target or indirect inhibition or activation of one or more other relevant targets. Drug-target binding, in turn, depends on [free drug in the target compartment](t) and [free target binding site](t), the former of which depends for oral drugs on the NLD balance between drug absorption (the source), clearance/metabolism (the permanent sink), and the fraction bound to a wide range of transient sinks, including off-target(s), membranes, plasma proteins, and lysosomes (Figure 3A), and the latter of which depends on the NLD balance between the target and overall system (noting that the same paradigm applies to off-target binding and the toxicodynamic responses thereto). Steady state free drug levels may be achieved, depending on the relative rates of the above processes vis-à-vis the dosing frequency.

**Figure 3.**
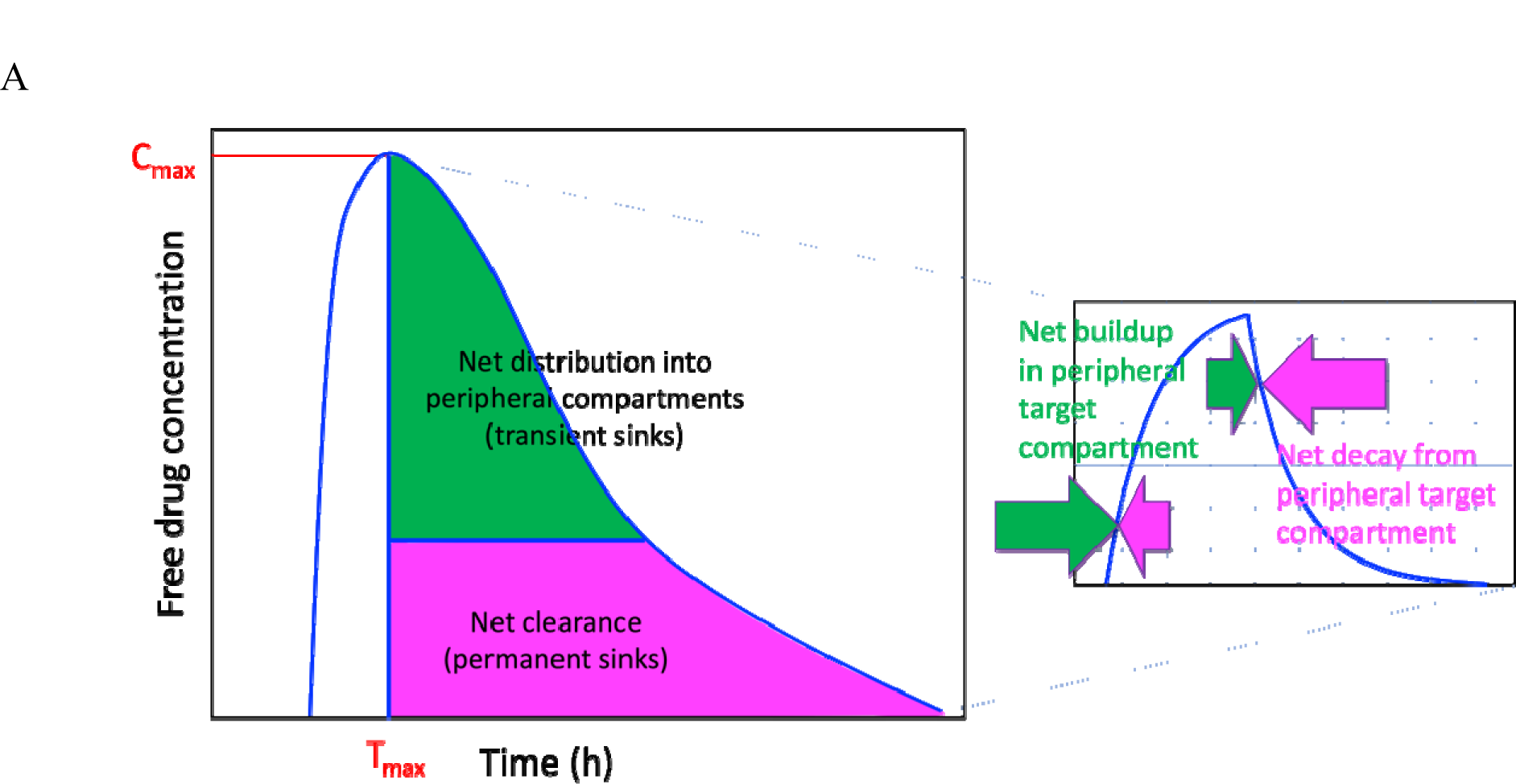
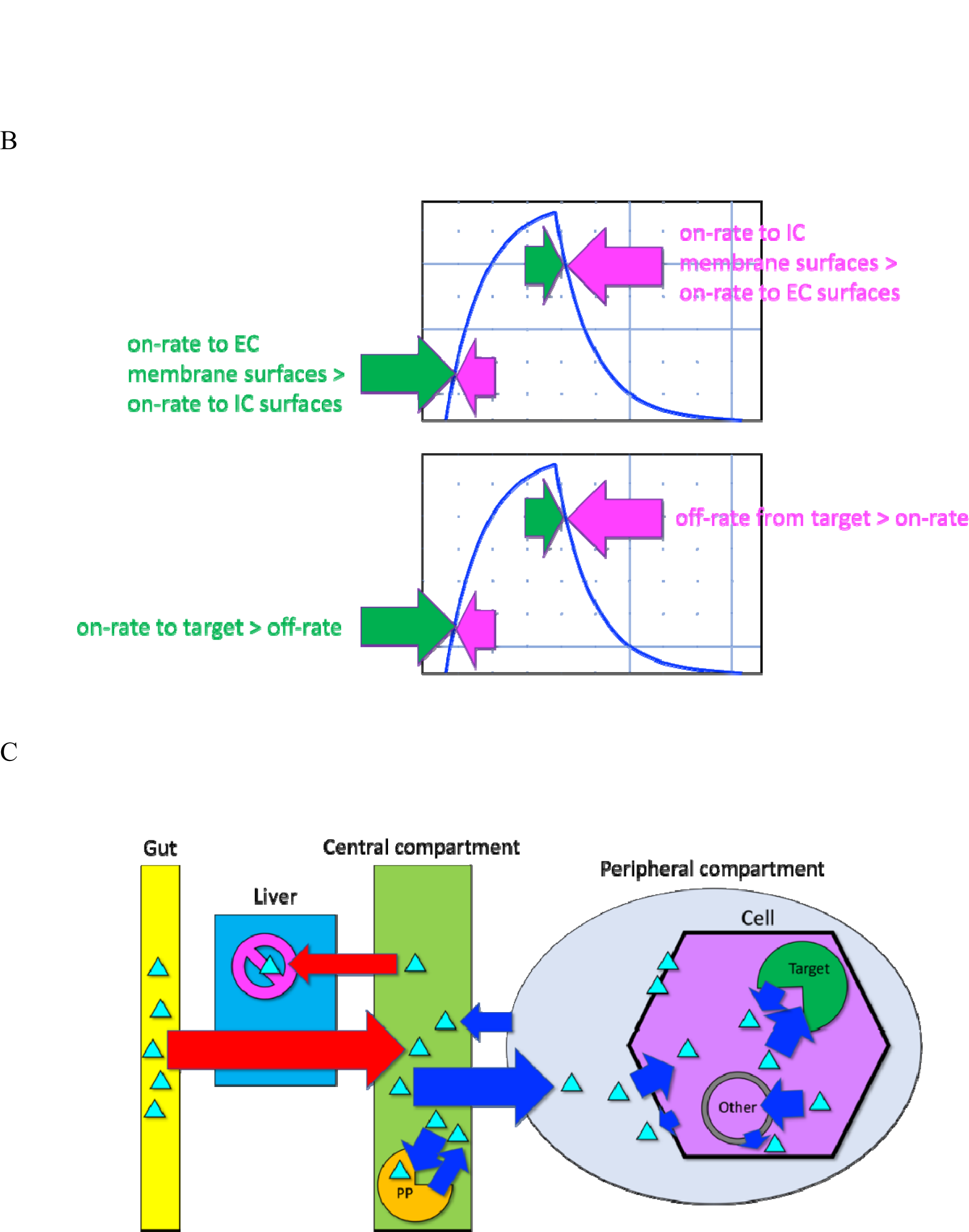
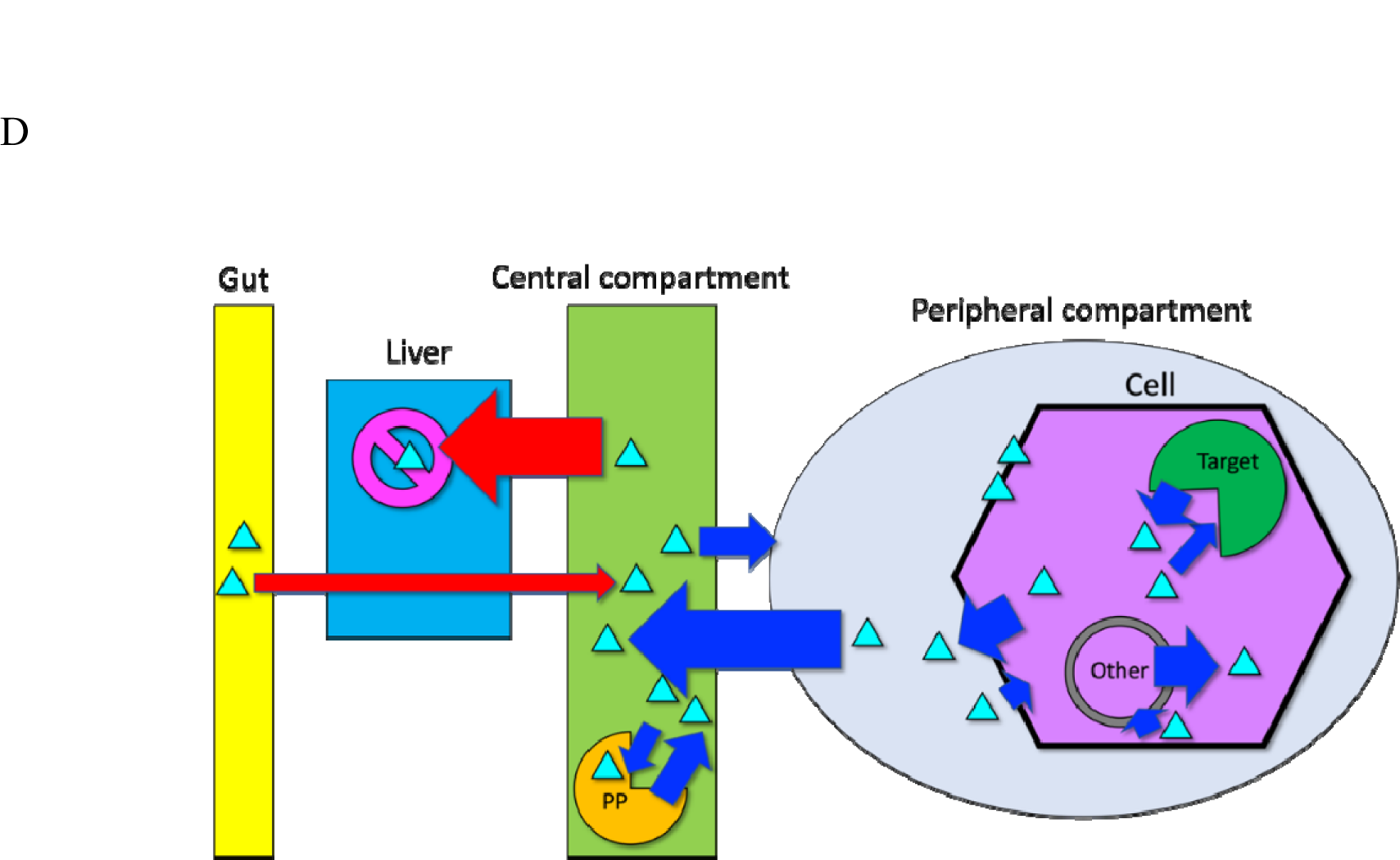
Drugs (cyan triangles) flow to and from the gut, central, and peripheral compartments along their concentration gradients in a highly non-linear source ➔ sink fashion. (A) Hypothetical single oral dose PK curve, in which a drug is absorbed from the gut, followed by first-pass metabolism, distribution of the remaining fraction into the peripheral compartments, and binding to a wide range of intra- and extracellular species (including plasma proteins). All of these processes operate concurrently, with faster absorption than distribution in the absorption (pre-C_max_) phase, faster distribution than clearance in the distribution (early post-C_max_) phase, and faster clearance than distribution in the clearance (late post-C_max_) phase (resulting in reversal of drug flow back to the central compartment). (B) All source ➔ sink processes (including transient distribution to/from peripheral compartments and binding, as well as permanent metabolic sinks) are bidirectional, with the fastest direction at a given time predominating (noting that lags in this process are introduced by slow off-rates from transient sinks). (C) The rates of drug entry into the central and peripheral compartments (red and blue arrows, respectively) are governed by the in-rate/out-rate balance to/from each compartment, where in-rate exceeds out-rate during the absorption phase. The fraction of drug bound to the various endogenous species residing within each compartment builds when the partner-specific on-rate exceeds the off-rate. (D) The rates of drug exit from the central and peripheral compartments (red and blue arrows, respectively) are governed by the out-rate/in-rate balance to/from each compartment, where the out-rate exceeds the in-rate during the clearance phase. The fraction of drug bound to its various endogenous partners decays when the partner-specific off-rate > on-rate.

The total free drug concentration in an in vivo system is given symbolically by:

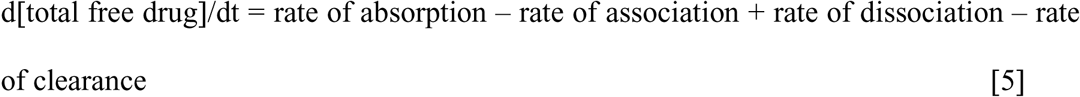

where, for the sake of simplicity, the association and dissociation terms represent the aggregate of all endogenous species to which the drug binds. The direction of drug flow through the system is determined by the in-rates to the various compartments and on-rates to the various endogenous species within. Inward flows to the peripheral compartments during the absorption/distribution phases are driven by the higher drug concentrations outside of those compartments (i.e., where the in-rates exceed the out-rates) (Figures 3B and 3C), which reverses during the clearance phase (i.e., where the out-rates exceed the in-rates) (Figures 3B and 3D). Lags in this process are introduced by slow off-rates from one or more transient sinks (including drug-target in cases of slow k_off_).

The <u>total</u> free drug concentration in the central compartment, representing the fraction of total drug available for distribution, binding, and clearance, is given symbolically by:

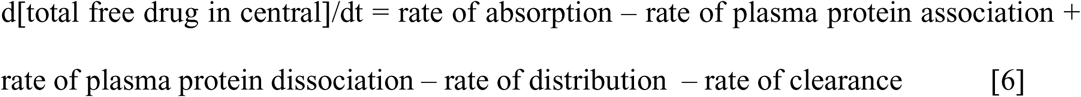

noting that the extremely high plasma protein (PP) concentration (∼500-700 μM (Merlot et al., 2014)) results in fast drug-PP on-rates:

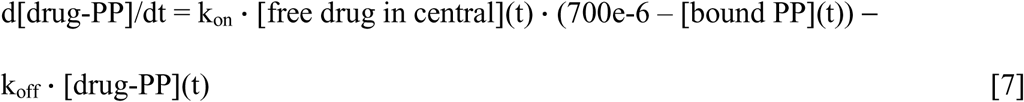

The free drug concentration in the extra-cellular compartment is given symbolically by:

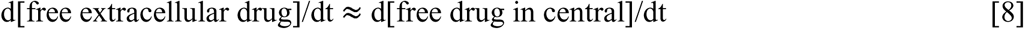

The free drug concentration in the intra-cellular target compartment is given symbolically by:

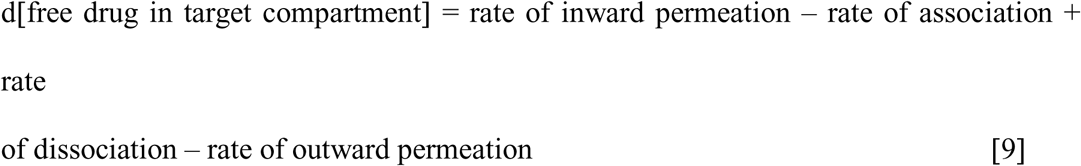

where the rate of inward permeation depends on [free extracellular drug](t), together with the rates of membrane entry and exit, and the rate of outward permeation depends on [free drug in target compartment](t), together with the rates of membrane entry and exit (noting that slow membrane exiting in either direction results in membrane accumulation/partitioning).

The optimal PK-PD and PK-TD scenarios may now be described as follows (under the tacit assumption that PD is governed directly by the time-dependent drug-bound target fraction, and TD is governed directly by the time-dependent drug-bound fraction of the most toxic off-target):

1. Kinetically tuned <u>drug-target</u> binding, resulting in the minimum γ_eff_ at the lowest possible free C_max_.
2. Kinetically mistuned <u>drug-off-target</u> binding, resulting in minimum off-target occupancy at the therapeutic free C_max_.
3. Slow drug-PP k_on_/fast k_off_, such that [drug-PP](t) << rate of drug distribution.
4. [total free drug](t) > rate of clearance.
5. Clearance of the drug-target complex (-[drug-target](t)) < the rate needed to achieve the minimum γ_eff_.
6. n · K_d_ < [free drug in target compartment](t), where n · K_d_ is the minimum efficacious exposure in the <u>kinetically tuned</u> binding scenario (e.g., 19 · K_d_ ➔ 95% drug-target occupancy). [free drug in target compartment](t) necessarily overshoots and decays back to the minimum efficacious exposure during the dosing interval (Figure 4A). Inter-dose troughs in [drug-target](t) result when [free drug in target compartment](t) decays to << n K_d_.
7. Dosing interval/quantity is commensurate with conditions 3-5.
8. [total free drug](t) < upper safe limit (allowing for dose escalation due to overdose or drug-drug-interactions (DDIs)) (Figure 4B).

**Figure 4.**
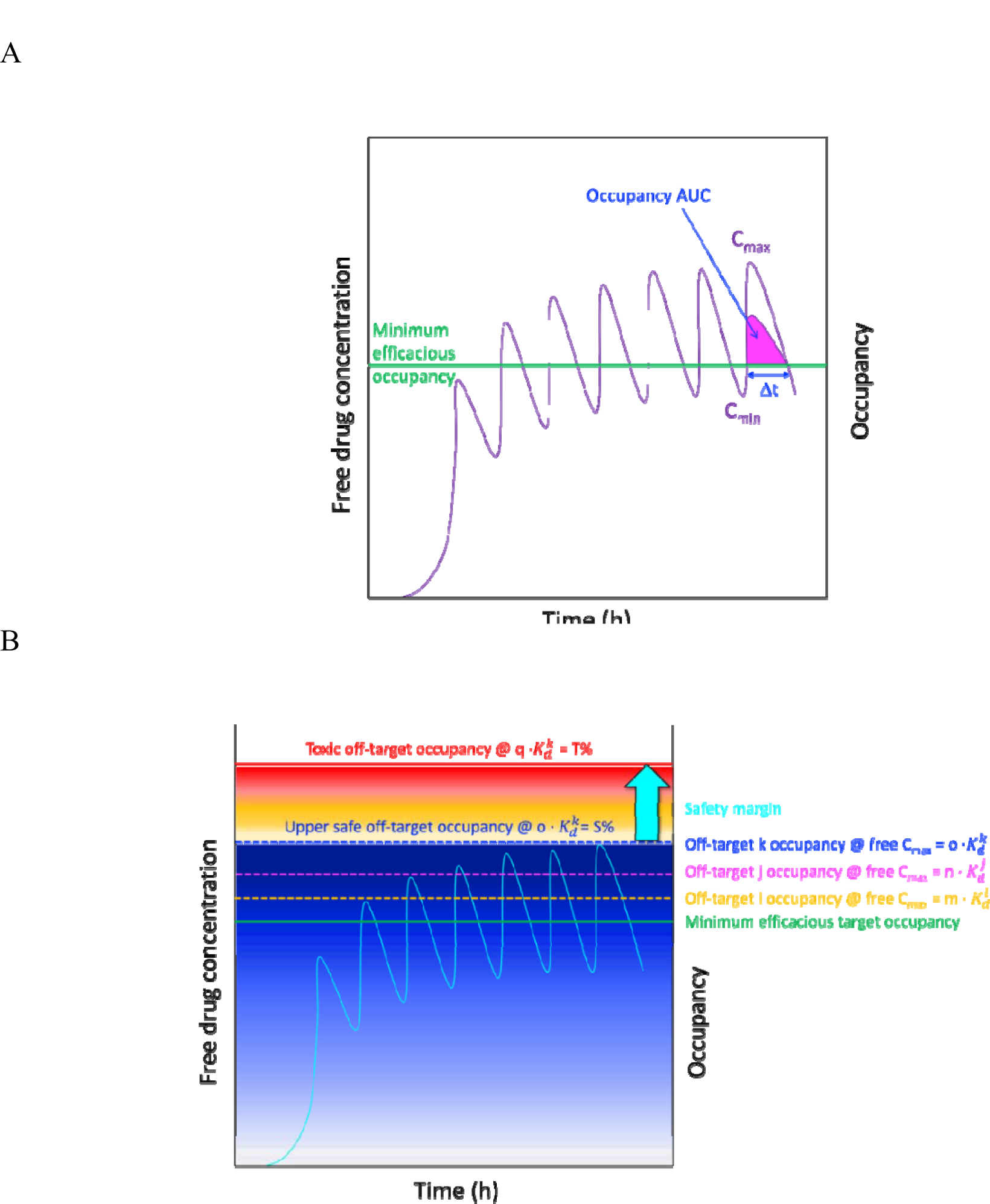
(A) Hypothetical multi-dose plasma PK curve, in which the free drug exposure builds to and overshoots the minimum γ_eff_ (topping out at the C_max_), followed by decay to the sub-efficacious level (bottoming out at the C_min_). The duration of action per dose (Δt) is governed by the degree to which the minimum γ_eff_ is overshot (represented by the area under the occupancy curve (AUC)). The minimum γ_eff_ is achieved at the lowest possible free C_max_ in cases of kinetically tuned binding. (B) Occupancy of a set of hypothetical off-targets i, j, and k as a function of the free plasma drug concentration ranging from the minimum efficacious exposure to the free C_max_, where the 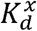 are the off-target binding constants (assuming the worst-case scenario of <u>kinetically tuned</u> off-target binding), n 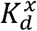 = C_max_ in multiples of 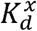, and m · 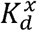 = the toxic free drug concentration for off-target k in multiples of 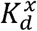.

The <u>mutual</u> effects of drugs on in vivo systems (i.e., PD) and in vivo systems on drugs (i.e., PK) is well appreciated in pharmaceutical science. The need for tight integration in PK-PD models (perhaps more than is commonly appreciated) follows from the overtly NLD relationship among the underlying behaviors shown in Figure 3. Nevertheless, PK, potency, and efficacy are routinely characterized separately and later reassembled into PK-PD models, representing tacit linearization of these highly non-linear processes. Equilibrium relationships between free drug concentration and occupancy are widely assumed in such models (equation 4), despite the known time dependence of the total drug concentration and frequent time dependence of the targeted binding site concentration (in which case, kinetically mistuned drug-target binding resulting in reduced P(CC | L, H) and P(CTI | CC, L, H) is conceivable) (Figure 5). This practice may result in drug failures when the equilibrium γ measured under in vitro conditions exceeds the steady state γ_eff_ under in vivo conditions. Furthermore, lead optimization is hampered by poor understanding of structure-ADME, structure-target binding, and structure-off-target binding relationships that may likewise contribute to drug failures.

**Figure 5.**
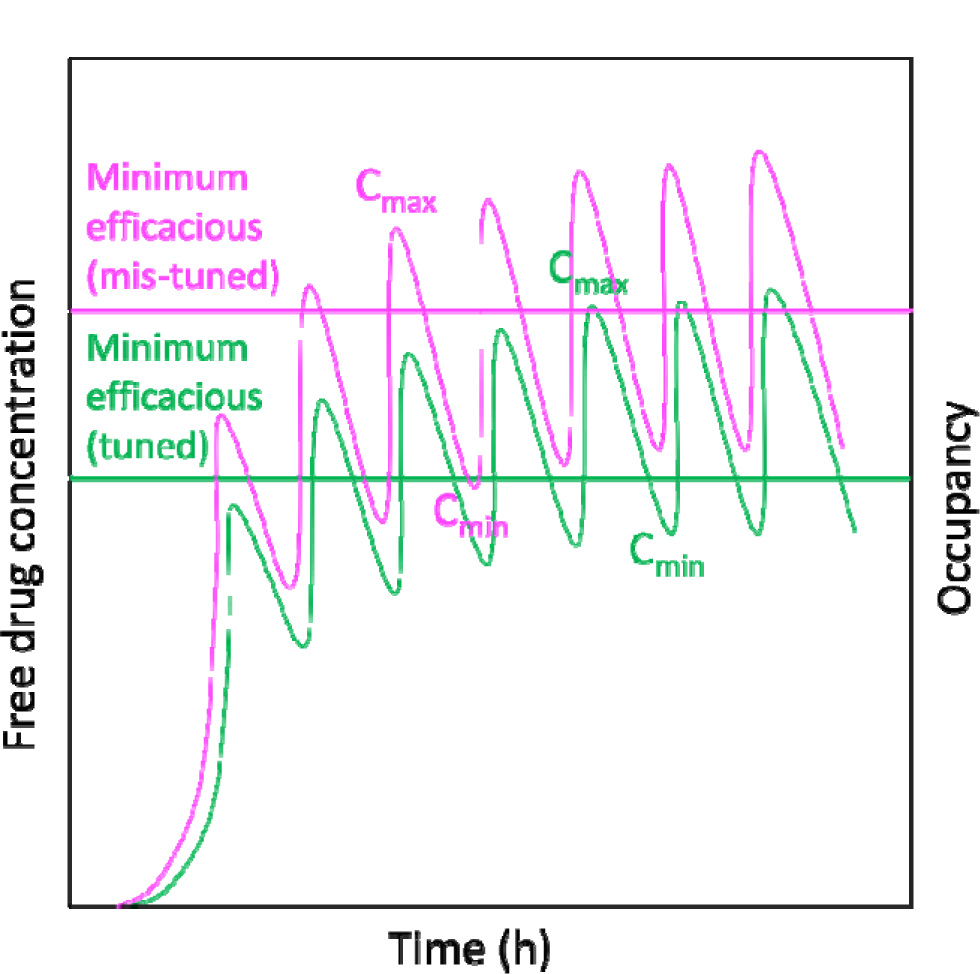
Kinetically mistuned drug-target binding results in a higher-than-expected minimum efficacious drug exposure in the target compartment under in vivo compared with in vitro conditions (in proportion to the gap between the rate of drug or target/binding site buildup and k_on_), necessitating higher in vivo drug exposures to achieve the γ_eff_ (Selvaggio and Pearlstein, 2018). The seamless connections between PK, binding site dynamics, binding kinetics (drug and target and off-target desolvation and resolvation costs) and dynamic fractional occupancy, and between dynamic fractional occupancy and disruption of one or more Yin-Yang balances underlying toxicity (TD) or restoration of a disease-causing Yin-Yang imbalance (PD) are summarized in Figure 6.

**Figure 6.**
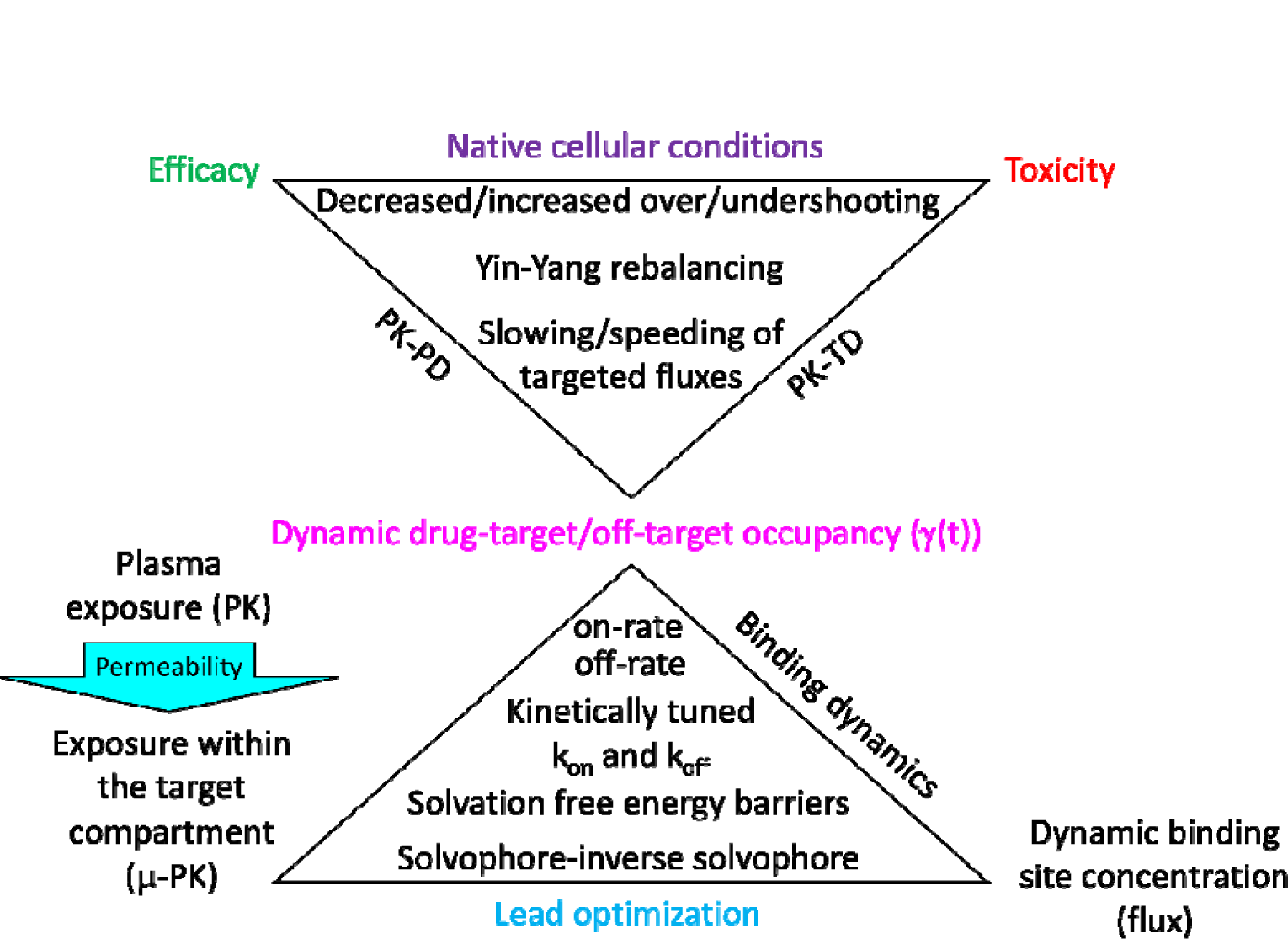
Dynamic fractional drug-target and drug-off-target occupancy builds and decays over time in step with buildup of the free drug concentration in the target and off-target compartments, buildup of the free target and off-target binding sites and decay of the free binding sites. k_on_ and k_off_ are governed by the desolvation and resolvation costs of drug and target/off-target binding sites. Afflicted Yin-Yang imbalances underlying a given cellular disease are corrected by dynamic fractional drug-target occupancy, whereas dynamic fractional off-target occupancy causes Yin-Yang imbalances underlying adverse effects.

### The implications of our proposed solvation free energy model for medicinal chemistry, computational chemistry, structural biology, and data science

Binding sites are classically viewed as molecular surfaces capable of forming complementary interatomic contacts with cognate partners, and non-covalent intra- and intermolecular free energies are widely assumed to originate principally from van der Waals, electrostatic, H-bond, π-π, π-cation, hydrophobic, hydrophilic, lipophilic and other interatomic contributions (the basi of force-fields, docking scoring functions, and molecular descriptors used for generating a wide variety of empirical/QSAR models). This hypothesis is predicated on the assumption that free energy contributions are similar for gases, solids, and aqueous solutes, which is inconsistent with the following observations:

1. Interatomic contact free energy contributions are far weaker than water-water and water-solute H-bonds under aqueous conditions (Young et al., 2007b; Pearlstein et al., 2010, 2017, 2021; Velez-Vega et al., 2015; Wan et al., 2020a, 2021a).
2. The search for complementary contacts is limited to short distances (close to the contact distance) due to the short-range nature of enthalpic interatomic contact contributions, as exemplified by the steep distance dependence of van der Waals and electrostatic energy contributions (1/r^6^ and 1/r^2^, respectively). Furthermore, the screening of electrostatic interactions by intervening water (due to the large dielectric constant of water) results in near zero enthalpic contributions at distances greater than the Debye length. As such, the means by which complementary contacts co-localize becomes a chicken-egg problem, commonly known in the protein folding arena as the Levinthal paradox (Karplus, 1997).
3. The magnitude of van der Waals enthalpy summed over numerous contacts in LMW and HMW molecules is negligible relative to the H-bond enthalpy of liquid water.
4. Enthalpic intra- and inter-solute H-bond and electrostatic gains are offset to varying degrees by the high desolvation costs of polar and charged partners (a zero-sum game at best).
5. π-π, π-cation, and halogen bonds are far weaker than water H-bonds. Under aqueous conditions, π-cation contacts are offset by the high desolvation costs of charged groups, and π-π contact and halogen bond free energy contributions may be mistaken for depletion of the H-bond enthalpy of water solvating aromatic and halogen groups (noting that halogens do not share their electrons, despite their high electronegativity).
6. The concept of hydrophobic (polar-non-polar), hydrophilic (polar-polar), and lipophilic (i.e., non-polar-non-polar) free energy contributions is strongly questioned by the lower enthalpies of water-methane compared with methane-methane interactions (which are significantly higher than that of water-water interactions) calculated using *ab initio* quantum mechanics (Pearlstein et al., 2017).
7. The inability of interatomic contacts to explain free energy barriers, which by definition, are comprised of separate unfavorable contributions in both the association/entry and dissociation/exit directions. Such contributions are necessarily graded and highly specific for cognate binding partners and functional intramolecular rearrangements, whereas unfavorable interatomic contact contributions are limited to all-or-none steric clashing and electrostatic repulsion that are indistinguishable in the association/entry and dissociation/exit directions.

Conversely, as we have demonstrated throughout this and our previous work, the H-bond free energy of solvating water is ideally suited for powering all non-covalent intra- and intermolecular rearrangements under aqueous conditions for the following reasons:

1. Water H-bond enthalpy is extremely large and highly sensitive to H-bond losses and gains in the presence of solutes (Young et al., 2007b; Pearlstein et al., 2010, 2017, 2021; Velez-Vega et al., 2015; Wan et al., 2020a, 2021a). Solutes are driven into structures in which the H-bond free energy losses and gains of their solvating water relative to bulk solvent are minimized and maximized, respectively. Structural rearrangements are thus powered by the medium (which stores and releases the preponderance of the available free energy in the presence of solutes) rather than interatomic contacts within and between the solutes themselves. This is reminiscent of the manner in which unfavorable potential energy is stored in space-time (the medium) in the presence of mass, which is released by driving the masses into their collapsed state (the true basis for gravity).
2. The non-polar regions of LMW and HMW solutes are solvated by high energy H-bond depleted water (relative to bulk solvent), the amount, magnitude, and spatial distribution of which are highly sensitive to the chemical structures and monomer sequences of LMW and HMW solutes, respectively.
3. H-bond depleted solvation is maximally present in the initial dissolved state (at t = 0) and is partially or fully expelled to bulk solvent over time via intra- and/or intermolecular rearrangements (folding, binding) to or toward the equilibrium distribution at fixed and variable concentrations, respectively (Figure 7A).
4. <u>Residual</u> H-bond depleted solvation that is non-auto-desolvatable in cis is invariably present to one degree or another in <u>post-rearranged</u> solute states, as follows:

a. At non-polar regions of external surfaces, including rearrangement interfaces.
b. In the case of HMW solutes, within internal/buried sub-surface cavities from which exchanges with bulk solvent are either slowed/impeded or abolished in the absence of external surface openings (Wan et al., 2020a, 2021a; Wan and Pearlstein, 2022).

**Figure 7.**
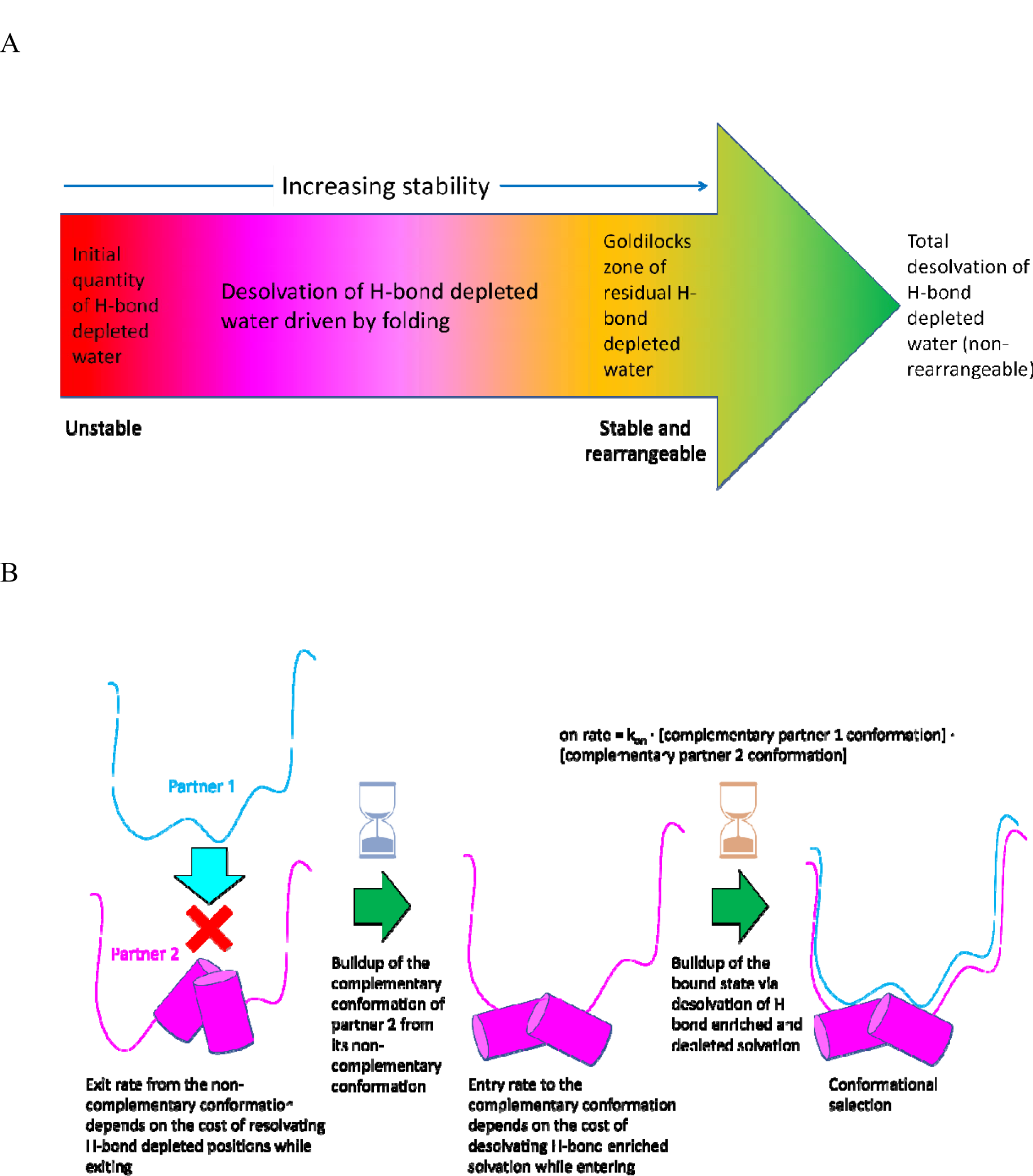
(A) The introduction of HMW and LMW solutes in water at t = 0 results in the generation of internal and external H-bond depleted solvation, the amount, magnitude, and distribution of which mirrors the non-polar surface composition. The minimum free energy states consist of those in which the H-bond <u>depleted</u> solvation has been maximally auto-desolvated via intramolecular rearrangement/folding and/or H-bond <u>enriched</u> solvation has been gained in the rearranged state. However, total auto-desolvation of such solvation results in non-rearrangeability of the populated states. (B) Non-covalent binding depends on both shape and solvophore-inverse solvophore complementarity. This is achieved via conformational selection rather than induced fit since the exit rates from non-complementary conformations and the entry rate to the complementary conformation are governed by resolvation in cis, independently of binding in trans.

Further intra- and/or intermolecular rearrangements initiated by perturbations in trans are powered principally by desolvation of this solvation (e.g., the introduction of a binding partner; change in membrane potential in the case of ion channels) (Wan et al., 2020a, 2021a; Wan and Pearlstein, 2022).

In addition, non-covalent free energy barriers and kinetic rate constants are well explained by unfavorable solvation free energy contributions, as follows:

1. Free energy barriers consist of varying proportions of enthalpic and entropic losses. However, entropic losses (as reflected in the state-dependent degree of unfavorable ordering of solvating water relative to bulk solvent and the loss of solute degrees of freedom in a given state) are non-specific. Furthermore, losses in solute entropy during association are always offset by varying degrees of entropic gains incurred from desolvation, and gains in solute entropy during dissociation are always offset by varying degrees of entropic losses incurred from resolvation of the dissociated partners.
2. k_on_ (the intermolecular association rate constant) and k_in_ (the intramolecular entry rate constant) are largely proportional to the mutual <u>desolvation</u> costs of the binding partners and rearrangement interface, respectively (Pearlstein et al., 2010). Desolvation costs, in turn, are proportional to the H-bond free energy of H-bond enriched solvating water (denoted as G_solv_) expelled from inter- or intramolecular rearrangement interfaces relative to bulk solvent (denoted as G_bulk_), where G_bulk_ − G_solv_ > 0. The lower the G_solv_ of this water, the higher the desolvation cost.
3. k_off_ (the intermolecular dissociation rate constant) and k_out_ (the rate of exit from a given intramolecular state) are proportional to the mutual <u>resolvation</u> costs of the binding partners and rearrangement interface, respectively (Pearlstein et al., 2010). Resolvation costs, in turn, are proportional to the H-bond free energy of solvating water (relative to bulk solvent) returning to H-bond depleted positions of the interface, where G_solv_ − G_bulk_ > 0. The higher the G_solv_ of this water, the higher the resolvation cost.
4. The <u>maximum</u> association/entry and dissociation/exit barriers are proportional to the number/magnitude of the H-bond free energy of H-bond enriched and H-bond depleted solvation, respectively.
5. Association/entry barriers can be lowered in a specific fashion via replacement of the H-bonds of expelled H-bond enriched water by intra- or inter-solute H-bonds during the rearrangement (such replacements are maximally present in cognate partners and functional intramolecular rearrangement interfaces and absent or present to lesser degrees in non-cognate partners and non-functional intramolecular rearrangement interfaces). The dissociation/exit barrier can be increased via the expulsion of graded amounts of H-bond depleted solvation, the unfavorable resolvation of which during dissociation/exiting is likewise graded (subserving a wide range of k_off_ and k_out_ magnitudes that are matched to the functional requirements at hand).

Lastly, conformational selection is favored by the solvation free energy model over induced fit binding, given that the formation of complementary binding partner conformations depends on:

1. Exiting one or more non-/less complementary conformations at rates governed principally by the cost of resolvating unfavorable H-bond depleted surface positions (in cis) (equating to the free energy gained via desolvation of H-bond depleted solvation while entering those conformations). As such, the persistence of non-complementary conformations cannot be short-circuited by desolvation of the associating partners in trans.
2. Entering the complementary conformations at rates determined principally by the desolvation cost incurred while entering.
3. A corollary to this is that intra- and intermolecular rearrangements (including the opening of so-called cryptic pockets) leading to the <u>de novo</u> generation of H-bond depleted solvation are slowed in proportion to the magnitude of the solvation free energy loss (i.e., the association or entry barrier is <u>increased</u> by the formation of such solvation, which largely negates the value of such pockets).

Binding is therefore limited to the following scenarios (Figure 7B):

1. Persistent (preorganized) complementary conformations of one or both partners, resulting from high resolvation costs incurred during exiting (translating to slow exit rates from such conformations).
2. Complementary conformations that form spontaneously (in cis) from one or more non-complementary conformations at sufficiently fast rates of entry to the former and exit from the latter (where “fast” is defined as non-rate limiting relative to the on-rate). This scenario may be incorrectly interpreted as induced fit. However, the complementary conformations are generated in cis rather than in trans.

The total non-covalent binding free energy can be derived as follows:

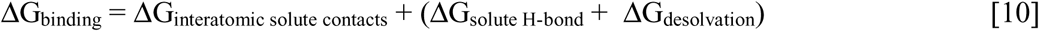

where ΔG_binding_ is the total free energy gain, ΔG_interatomic solute contacts_ is the interatomic contact free energy contribution, ΔG_desolvation_ is the water transfer free energy contribution, and ΔG_solute H-bond_ is the intra- or inter-solute H-bond free energy contribution. ΔG_interatomic solute contacts_ can be neglected to a first approximation, since (ΔG_solute H-bond_ + ΔG_desolvation_) >> ΔG_interatomic solute contacts_ (Pearlstein et al., 2017):

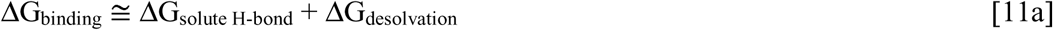

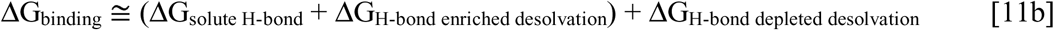

where:

a. ΔG_interatomic solute contacts_ and ΔG_solute H-bond_ are always < 0.
b. ΔG_H-bond enriched desolvation_ is always > 0.
c. ΔG_solute H-bond_ + ΔG_H-bond enriched desolvation_ ➔ 0 as ΔG_solute H-bond_ ➔ ΔG_H-bond enriched desolvation_.
d. ΔG_binding_ ➔ ΔG_H-bond depleted desolvation_ (given perfect replacement of the H-bonds of desolvated H-bond enriched solvation).

*Caveat 2: The interpretation of structure-activity relationships (the currency of medicinal chemistry) on the basis of interatomic contact free energy contributions, the prediction of non-covalent free energy and structure-free energy relationships using force-field based approaches (e.g., molecular dynamics, free energy perturbation, conformational searching, energy minimization), and data modeling approaches (e.g., quantitative structure-activity relationship (QSAR) analysis, docking/scoring, pharmacophore analysis) are subject to grossly overestimated solute and grossly underestimated solvation free energy contributions*.

Equation 11b can be expressed as:

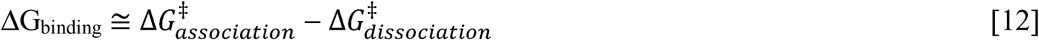

where 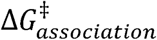 and 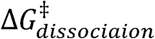 are the free energy barriers for entry and exit to/from a given intermolecular state, respectively. Structure-kinetics relationships can therefore be explained on the following basis (contrary to the classical structure-free energy interpretation):

1. The non-covalent entry free energy barrier magnitude equates to the net desolvation cost, consisting of the H-bond free energy lost in transferring H-bond enriched solvation to bulk solvent, offset by the free energy gained via intra- or inter-solute H-bond formation (typically > 0 and rarely < 0). As such, the populated non-covalent states are those entered the fastest, rather than those stabilized by intra- or inter-solute H-bonds and electrostatic contacts (as is widely assumed). The most frequently visited states are those in which the lowest desolvation cost of H-bond enriched solvation (which we refer to as “gatekeeper solvation”) is incurred during entry. A simple rule of thumb is that H-bonds observed in folded HMW species and complexes reflect the allowed positions of polar groups during folding or binding (i.e., where H-bond enriched solvation existed previously), whereas non-polar contacts reflect H-bond depleted positions that were desolvated during folding or binding.
2. The magnitude of the exit free energy barrier equates to the total <u>resolvation</u> cost incurred during the decay of a given non-covalent intra- or intermolecular state <u>at H-bond depleted positions</u>. Non-covalent states are kinetically stabilized principally by this contribution. As such, the most persistent states are those whose resolvation costs incurred at H-bond depleted solvation positions in the exited/dissociated state are the highest. Such solvation necessarily resides within a Goldilocks zone in which:

a. Anti-solubilizing and destabilizing effects are balanced against low energy gatekeeper solvation.
b. Non-specific intra- and intermolecular rearrangements that expel H-bond depleted solvation are minimized.
c. Rate-determining steps (lags) due to slow off-rates of endogenous species are minimized.
3. Non-covalent states build and decay over time, wherein buildup exceeds decay in the net buildup phase (i.e., on-rate is increasing relative to off-rate) and decay exceeds buildup in the net decay phase (i.e., on-rate is slowing relative to off-rate) (Figure 3B, bottom).

## Materials and methods

In our previous work, we used molecular dynamics simulations together with WATMD (available at https://github.com/Novartis/watMD) to predict the positions and qualitative magnitudes of H-bond enriched, depleted, and bulk-like solvation on the external and internal surfaces of COVID M^pro^ (Wan et al., 2020a). We used the same methods in this work to predict the solvation properties of the M^pro^ inhibitors nirmatrelvir and PF-00835231. We refer to these simulations as “solvation dynamics (SD) simulations” since they are focused on water exchanges to/from bulk solvent and solvation rather than solute rearrangements (noting that solute rearrangements predicted using force-field-based methods are highly unreliable due to the overestimation of interatomic contact energies and underestimation of solvation free energy). Our method is fully described in references 5, 7, and 10 and briefly summarized here:

1. Nirmatrelvir (PF-07321332) (Kneller et al., 2022) and PF-00835231 (Boras et al., 2020) were extracted from their crystallized M^pro^-bound structures (PDB code = 7SI9 (Kneller et al., 2022) and 6XHM (Boras et al., 2020), respectively). The structures were prepared as described in the above references.
2. The inhibitors were simulated in their fully restrained conformations using AMBER 20 (Case et al., 2022) PMEMD CUDA (GAFF and ff99sb force-fields) for 100 nanoseconds (ns) in a box of explicit TIP3P water molecules.
3. The fully unrestrained monomeric M^pro^ structure (PDB code = 2QCY (Shi et al., 2008)) was simulated in our previous work (Wan et al., 2020a) using AMBER 16.
4. The time-averaged hydrogen (H) and oxygen (O) occupancies within a stationary three-dimensional grid comprised of 1 Å^3^ voxels in which each solute structure was embedded were calculated over the last 40,000 frames (10 ns) of a 100 ns trajectory using WATMD V9 (Velez-Vega et al., 2014, 2015; Pearlstein et al., 2017) (noting that the solvation fully converges within the 90 ns timeframe prior to the water data collection stage).
5. The high and low occupancy voxel data was normalized to the bulk-like solvating water (i.e., the mean of the distribution) on a solute-by-solute basis, so as to achieve self-consistency (WATMD calculations can be considered as first principles for this reason). The high occupancy voxels were then scaled to the largest voxel in the entire dataset.
6. The voxels were then assigned to bulk-like, H-bond enriched, and H-bond depleted solvation states (noting that the results in all cases are distributed in a Gaussian fashion, the specific properties of which vary among solutes), as follows:

a. **Bulk and bulk-like voxels** reside at the mean of the distribution, where the H and O positions in the same voxel are fully uncorrelated (corresponding to no orientational preference of the occupying water molecule, comparable to bulk solvent).
b. **H-bond depleted voxels** reside in the extreme left tail of the distribution (where G_bulk_ < G_solvation_), resulting from diminished solute H-bond partners compared with bulk solvent.
c. **H-bond enriched voxels** reside in the extreme right tail of the distribution (where G_bulk_ > G_solvation_), resulting from enhanced solute H-bond partners compared with bulk solvent.
7. Water occupancies are underestimated in voxels that are transiently occupied by mobile solute atoms that compete with water exchanges. This problem is circumvented as follows:

a. LMW solutes are simulated with full restraints (ideally in their crystallized bound conformations).
b. HMW solutes that undergo large rearrangements are simulated with light restraints (noting that water exchanges to/from tight spaces are artificially slowed by restrained solute motions, resulting in artificially magnified voxel sizes).
c. HMW solutes that undergo small, localized rearrangements (e.g., the β-hairpin in COVID M^pro^) are simulated without restraints, and each of the mobile region(s) are separately aligned across all of the frames during the WATMD calculations (and are therefore stationary relative to the voxel grid).
8. Overlays between the structures and predicted voxel occupancy data were visualized using PyMol 2.0 (Schrodinger, LLC), with the occupied voxels denoted by spheres whose sizes are proportional to the predicted H and O occupancies, and whose colors denote the preference for O (red), H (blue), and neither (white).

In our previous work, we studied the catalytic cycle and solvation properties of COVID M^pro^ vis-à-vis a set of published inhibitors based on the aforementioned principles (Wan et al., 2020a). We deduced that:

1. M^pro^ binds to substrates and inhibitors in its monomeric form.
2. The catalytic site is activated by substrate binding, followed by dimerization.
3. Turnover is followed by product dissociation, deactivation, and dimer dissociation.

Here, we predict the solvation properties of two published M^pro^ inhibitors and compare the complementarity thereof to that of the catalytic pocket.

## Results

In our previous work, we used WATMD calculations to characterize the inverse solvophore of monomeric COVID M^pro^ (PDB code = 2QCY) and its complementarity to the distribution of polar and non-polar groups on nirmatrelvir extracted from the dimeric M^pro^ complex (PDB code = 7SI9) and an analog thereof (PF-0083521) extracted from the dimeric M^pro^ complex (PDB code = 6XHM) (Figure 8) (noting that our LMW WATMD capability was unavailable at that time).

**Figure 8.**
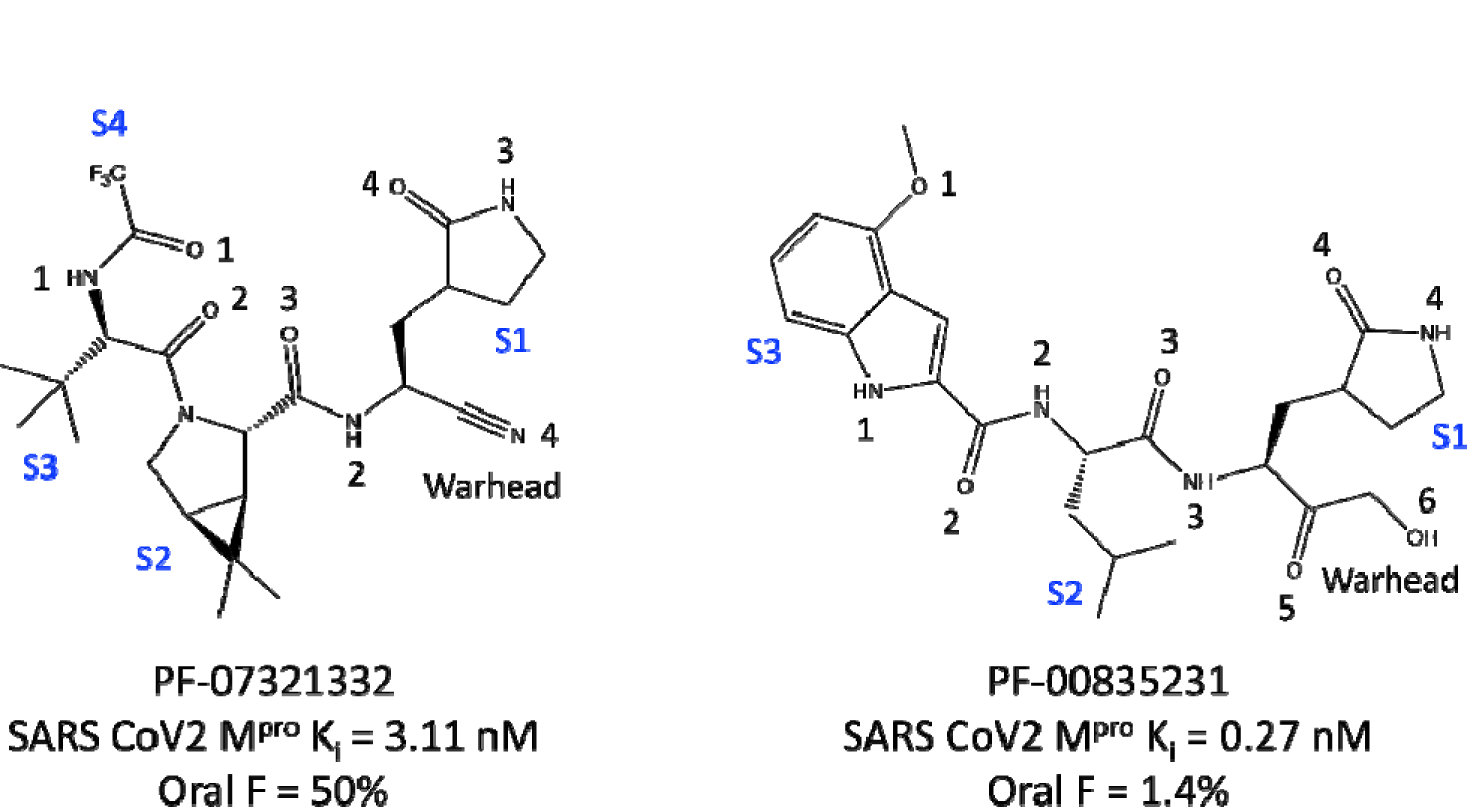
2D structures and experimental data for the COVID M^pro^ inhibitors PF-07321332 (nirmatrelvir) and a crystallized analog thereof (PF-00835231) (Boras et al., 2020; Owen et al., 2021). The inhibitor side chains are labeled S1-S4. Oxygen and nitrogen atoms are numbered consecutively in the N- to C-terminal direction.

Both PF-07321332 and PF-0083521 react covalently with the catalytic cysteine of M^pro^, resulting in accumulation of the covalent complex over time. The k_on_ of each inhibitor depends on th quality of mutual intermolecular drug-protein H-bond replacements for H-bond enriched water expelled during the association step (noting that energetically conserved replacements need not be spatially conserved). The rate of drug-target occupancy accumulation depends on the slower of the on-rate and covalent reaction rate relative to the drug-target off-rate (noting that efficacy depends on equal or faster buildup of the inhibitor-bound state compared with the rate of M^pro^-mediated polyprotein cleavage). It is reasonable to assume that the resolvation costs of the catalytic site and <u>native substrates</u> are evolutionarily calibrated for achieving k_-1_ k_cat_ (where k_-1_ is the substrate dissociation rate constant), whereas non-covalent inhibitors with smaller footprints that desolvate less H-bond depleted solvation than the native substrates likely exhibit k_off_ > k_-1_. It is likewise reasonable to assume that the H-bond <u>enriched</u> solvation in the catalytic site is organized for optimal replacement by the maximum common solvophore among all of the endogenous M^pro^ substrates of the polyprotein (Figure 9). The H-bond enriched and depleted solvation in the catalytic site is highly complementary to that of PF-07321332 and PF-0083521 (as described below). However, the actual k_on_, the pre-reacted non-covalent k_off_, and the post-reacted k_off_ cannot be quantitatively ascertained in the absence of measured binding kinetics data.

**Figure 9.**
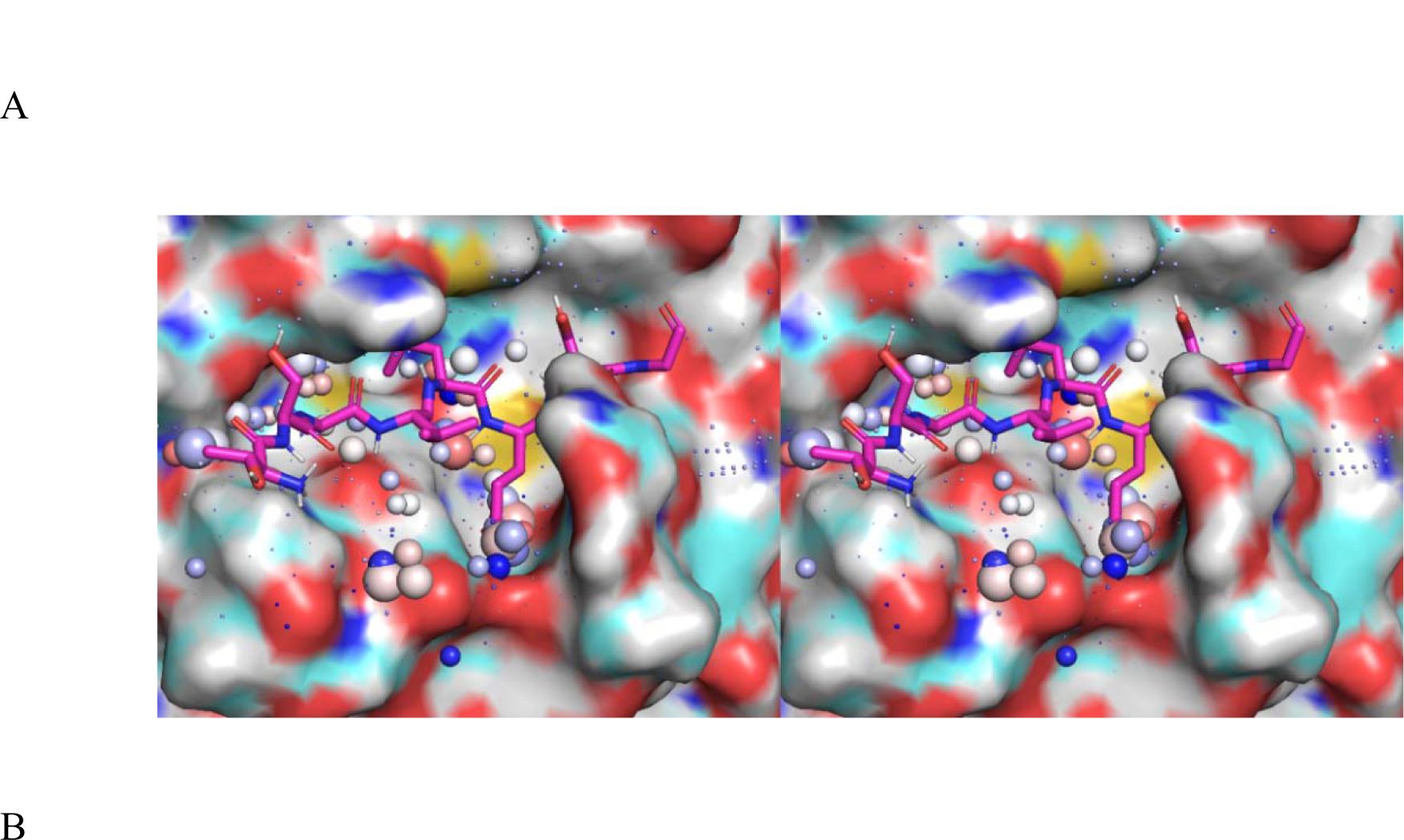
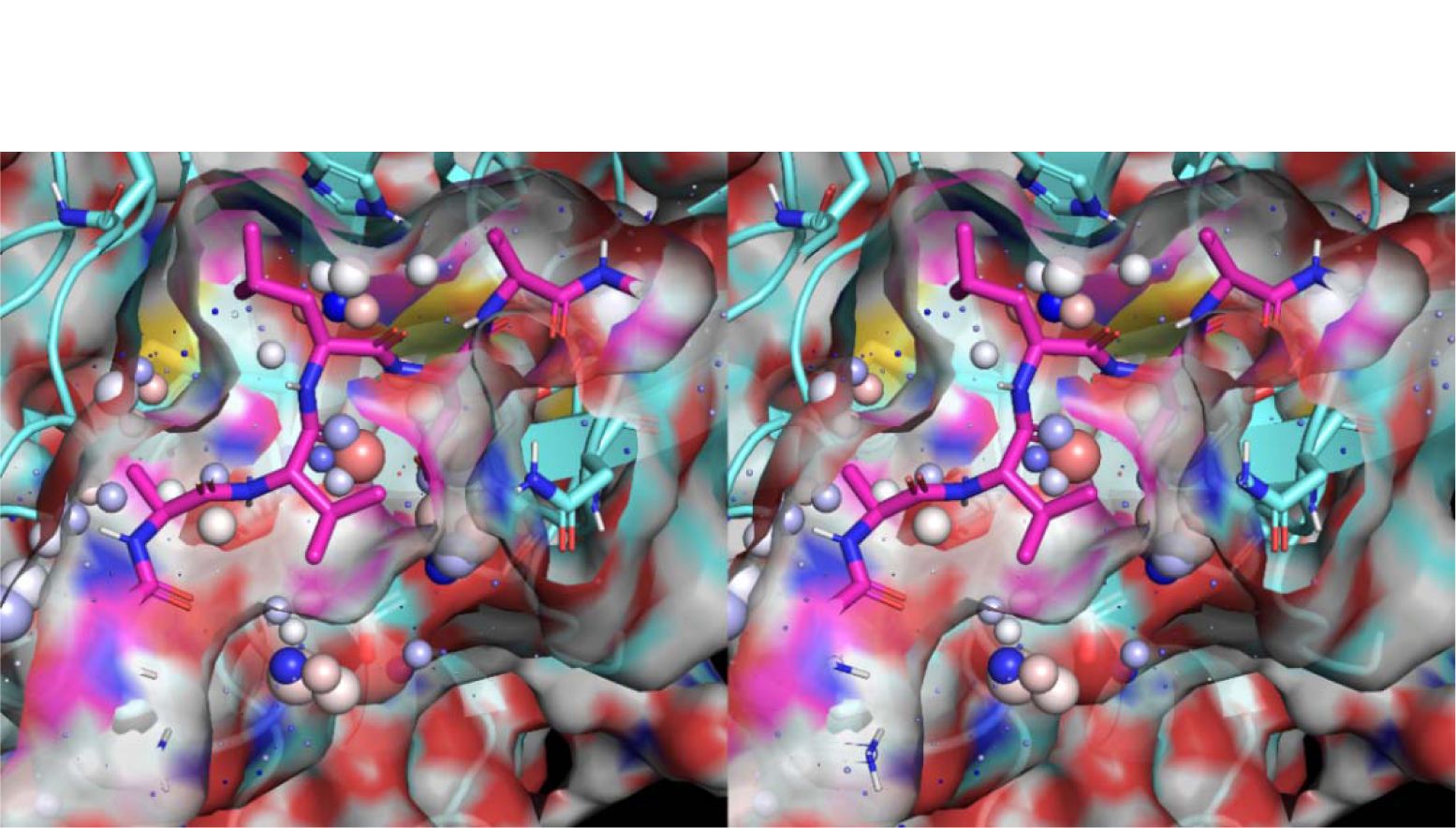
(A) Stereo view of the inverse solvophore of the apo monomeric mutant form of M^pro^ (PDB code = 2QCY) overlaid on an M^pro^ substrate extracted from PDB code = 2Q6G (Wan et al., 2020a). Low and high occupancy voxels are shown as color-coded spheres, as explained in Materials and methods (B) Same as A, except showing the solvent accessible surface of the substrate. All voxels contained within the substrate surface are expelled during association (noting that water is trapped within the northwestern region of the binding interface, and also exists in the southern region of the substrate-bound protein).

The solvophores of PF-0083521 (Figures 10A and 10B) and PF-07321332 (Figures 10C and 10D) share both similarities and differences in accordance with their unique shapes and polar/non-polar compositions and spatial distributions. Low occupancy/H-bond depleted voxels are distributed around the non-polar regions of both inhibitors, and high occupancy/H-bond enriched voxels are distributed as follows (referenced to the numbering shown in Figure 8):

**PF-0083521**: Clusters of high occupancy voxels are concentrated around HN2 (ultra-high occupancy), HN3, HN4, and both of the oxygen lone pair positions of C=O2. High occupancy voxels are distributed in a chain-like fashion within a surface groove containing C=O2 and HN3 bounded by the warhead and S2 moieties. Low-cost desolvation of this inhibitor depends on the quality of H-bond replacements for each solvating water by appropriate M^pro^ H-bond partner(s). Low-cost desolvation of M^pro^ likewise depends on the quality of H-bond replacements for each solvating water in the catalytic site by appropriate inhibitor H-bond partner(s).
**PF-07321332**: Clusters of high occupancy voxels are concentrated around HN1, HN2, and HN3 (with similar occupancy magnitudes at HN1 and HN2), together with both of the oxygen lone pair positions of C=O2. High occupancy voxels are distributed in a chain-like fashion within a surface groove containing HN2 (described in detail in Wan et al., 2021a). Desolvation of the high occupancy voxels at HN1 and HN2 is especially costly. Low-cost desolvation likewise depends on the aforementioned criteria.

**Figure 10.**
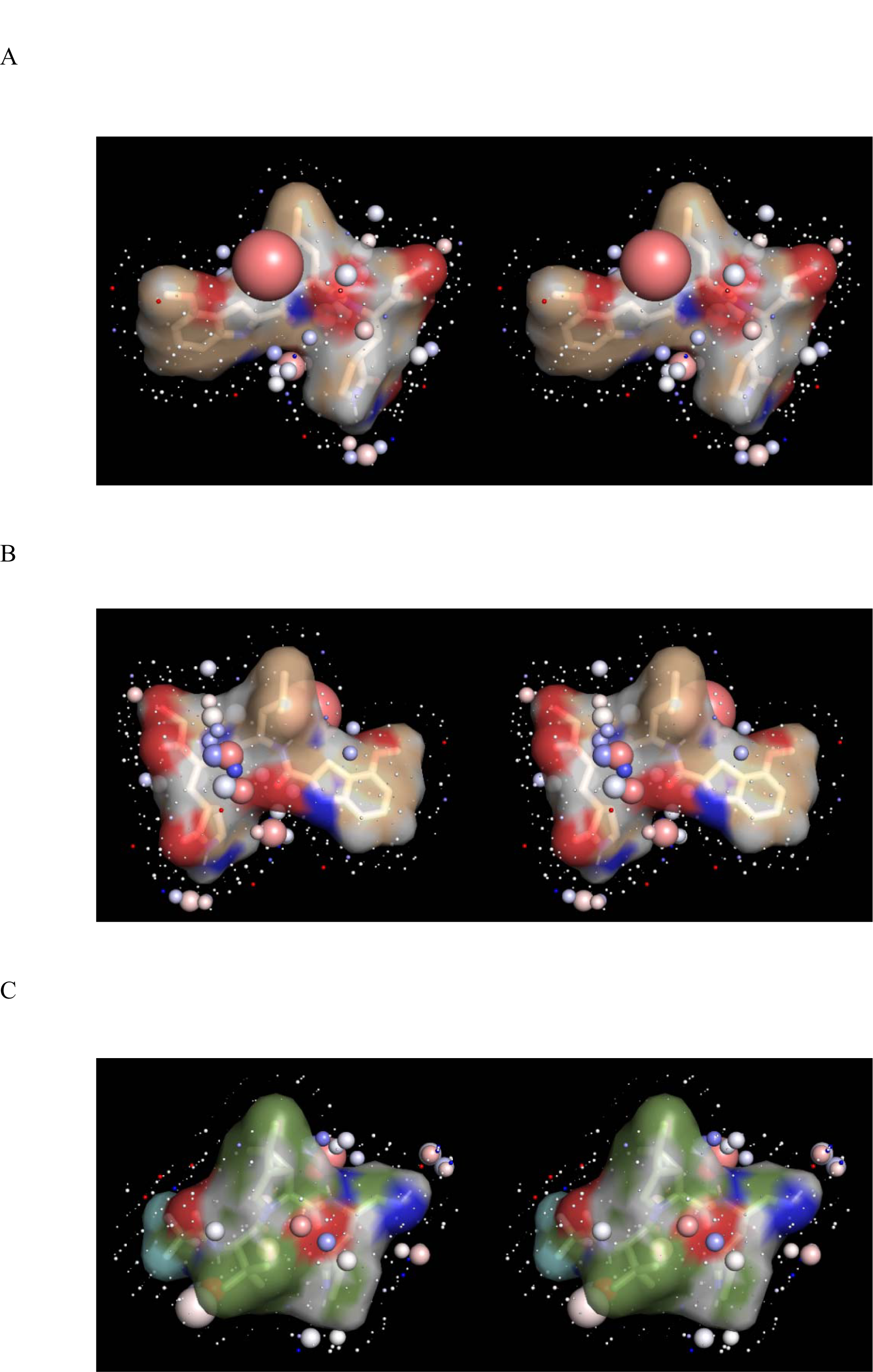
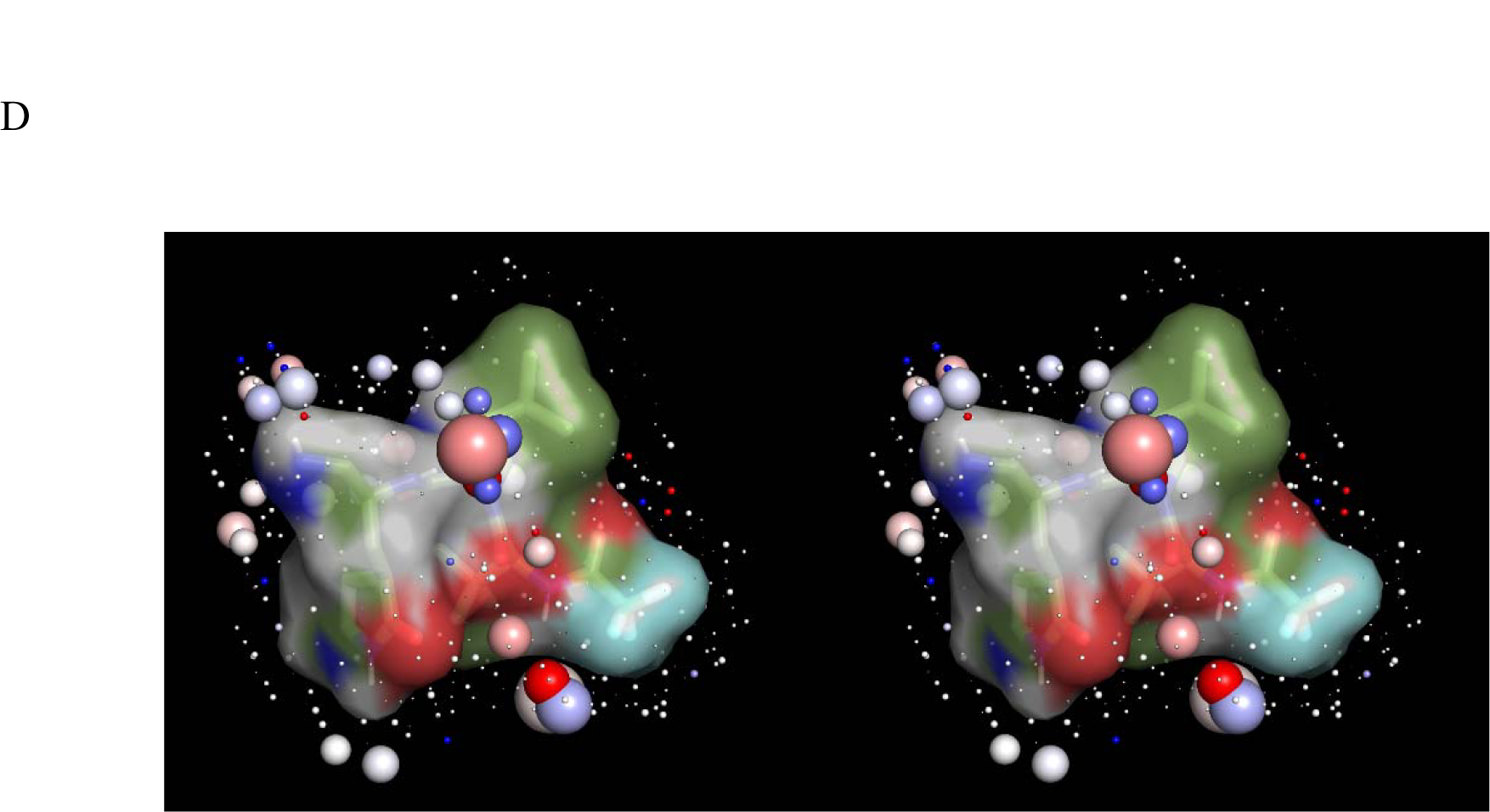
Stereo views of the structures and solvent accessible surfaces of the M^pro^ inhibitors PF-0083521 and PF-07321332 overlaid on their respective LMW solvophores calculated with WATMD (voxel annotations explained in Materials and methods). The inhibitor structures were extracted from the inhibitor bound M^pro^ crystal structures (see text), the warheads were manually restored to their pre-reacted states in preparation for the SD simulations, and the conformations were restrained in their crystallized conformations. (A) The solvent-facing side of PF-0083521 (tan sticks and transparent surface color-coded by atom type) with the reacted warhead on th right. (B) Same as A, except the pocket-facing side of PF-0083521 with the reacted warhead on the left. (C) The solvent-facing side of PF-07321332 (green sticks and transparent surface color-coded by atom type) with the reacted warhead on the right. (B) Same as C, except the pocket-facing side of PF-07321332 with the reacted warhead on the left.

The high potency of both PF-0083521 and PF-07321332 is consistent with good overall packing within the M^pro^-inhibitor interfaces (Figure 11) and high bidirectional complementarity between the polar groups and high occupancy/H-bond enriched voxels of M^pro^ and PF-0083521 (Figure 12) and M^pro^ and PF-07321332 (Figure 13). However, mismatches between the CF_3_ group of PF-07321332, high occupancy voxels in the P4 region, and suboptimal packing in the P2 region of M^pro^ (Figure 12A) may contribute to the ∼10-fold lower potency of this inhibitor (which is offset by its substantially higher bioavailability (Figure 8)).

**Figure 11.**
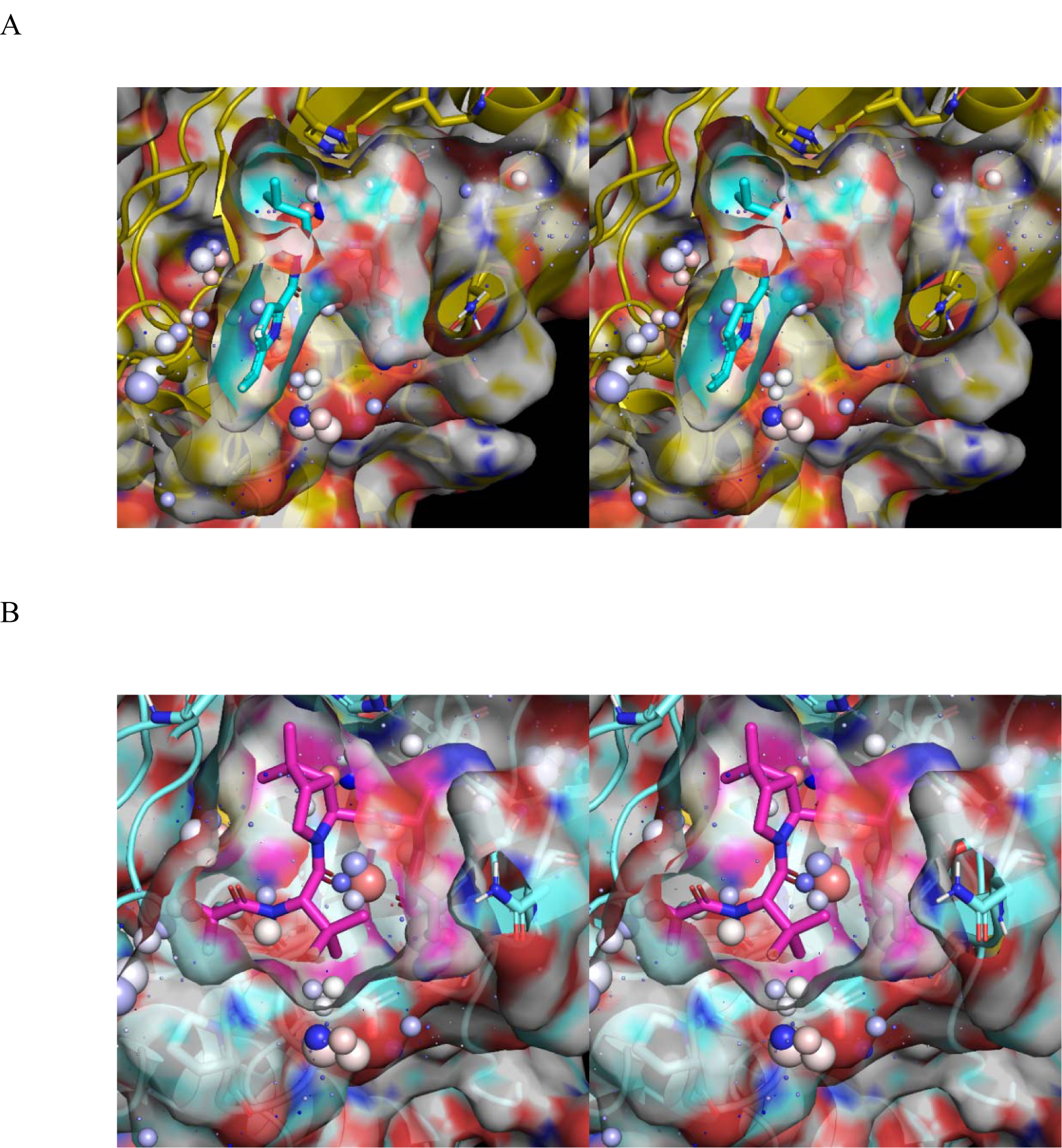
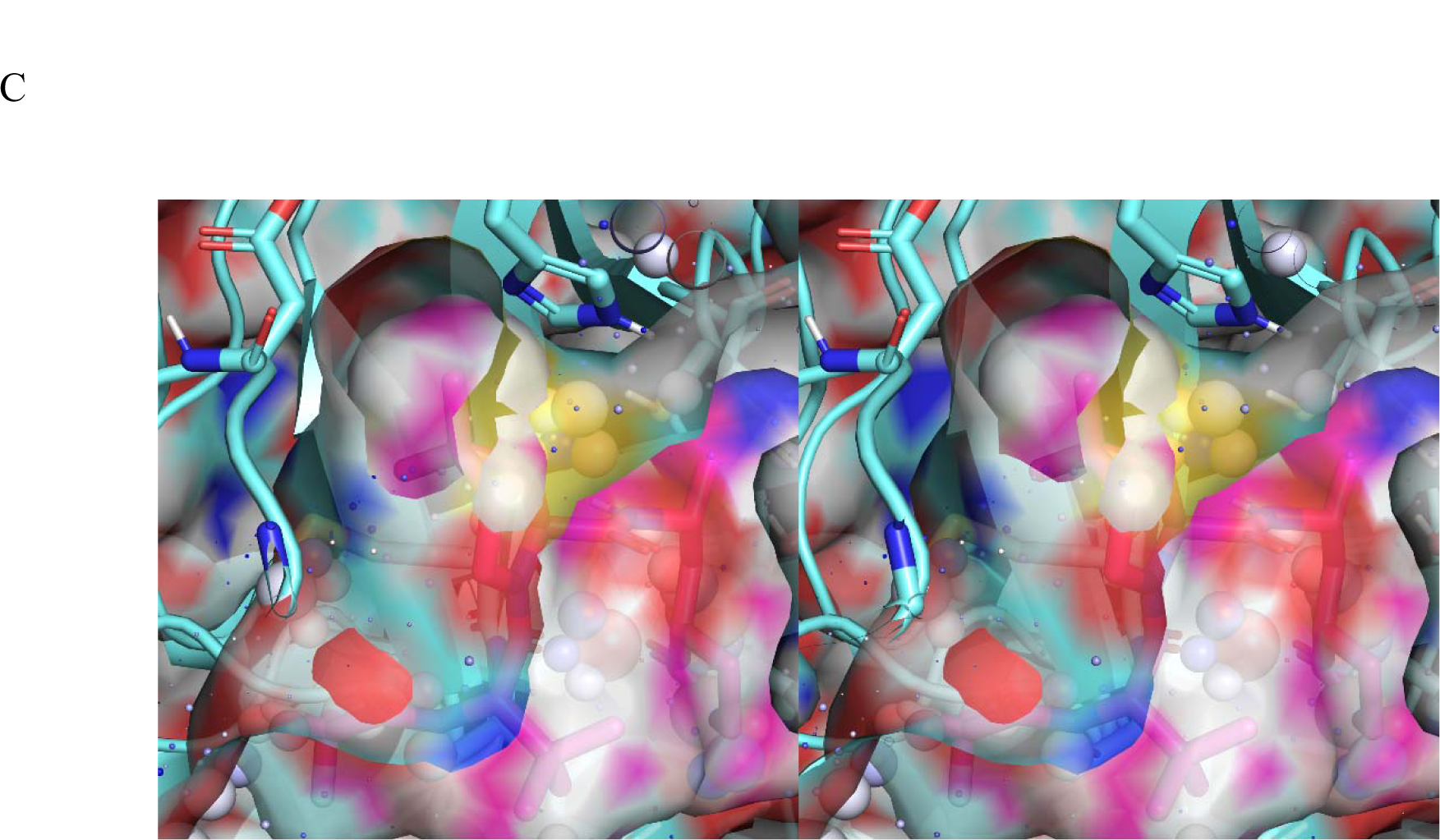
(A) Stereo view of the packing between apo monomeric M^pro^ (2QCY) and PF-07321332 (voxel annotations described in Materials and methods). All voxels contained within the substrate surface are expelled during association. (B) Same as A, except for PF-0083521 extracted from 6XHM. (C) Same as B, except showing the comparatively poor packing in the P2 pocket region occupied by trapped water.

**Figure 12.**
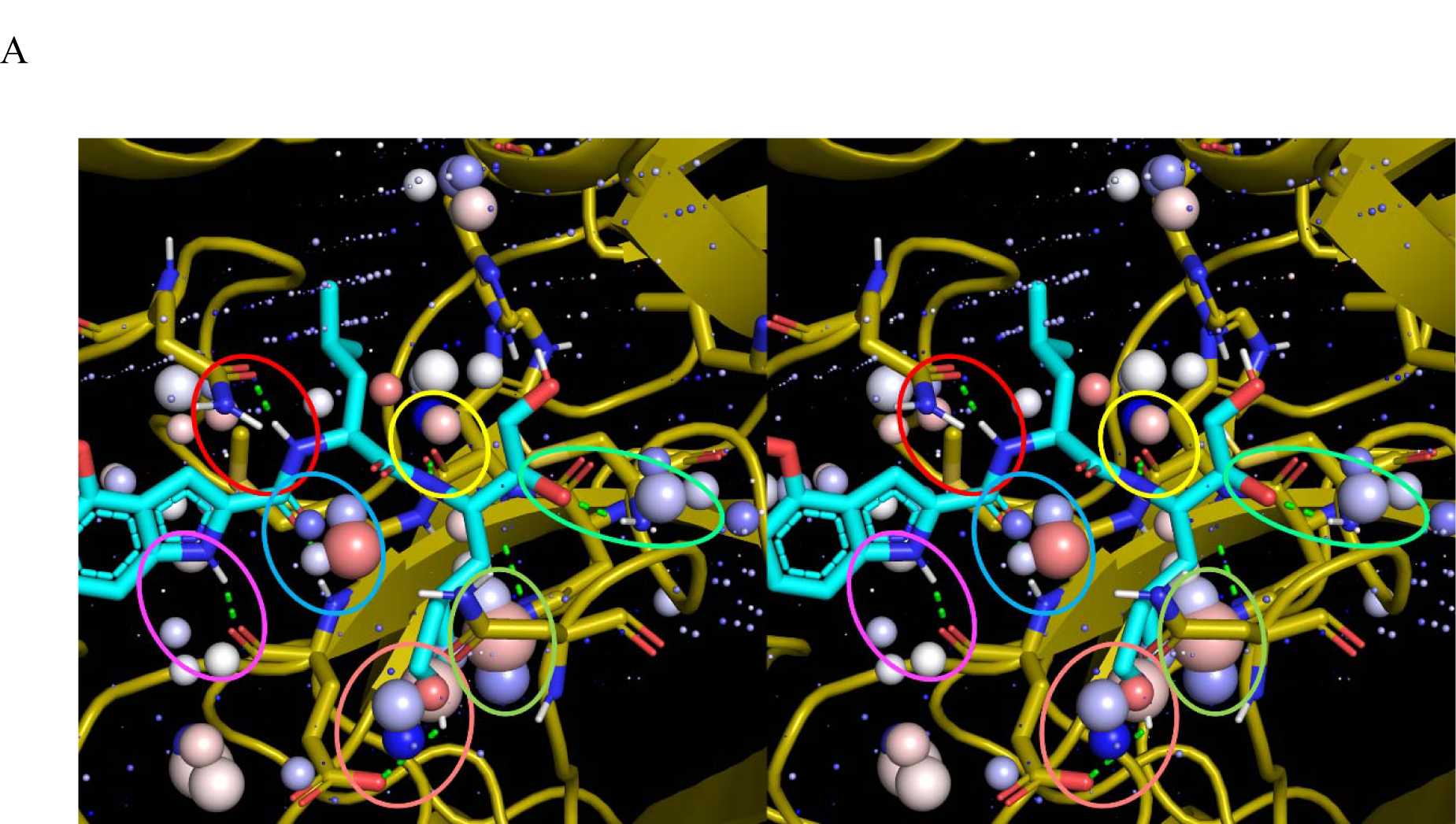
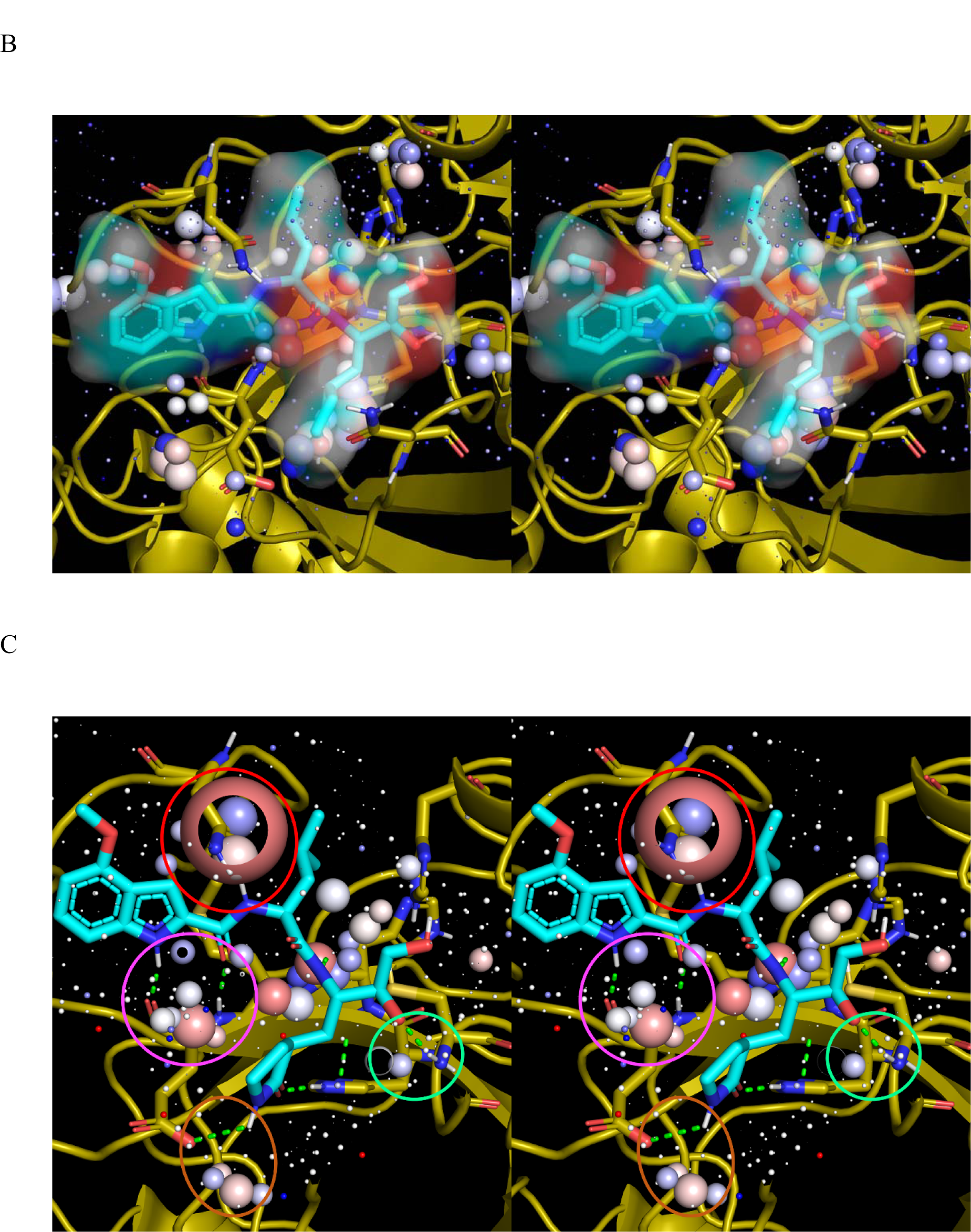
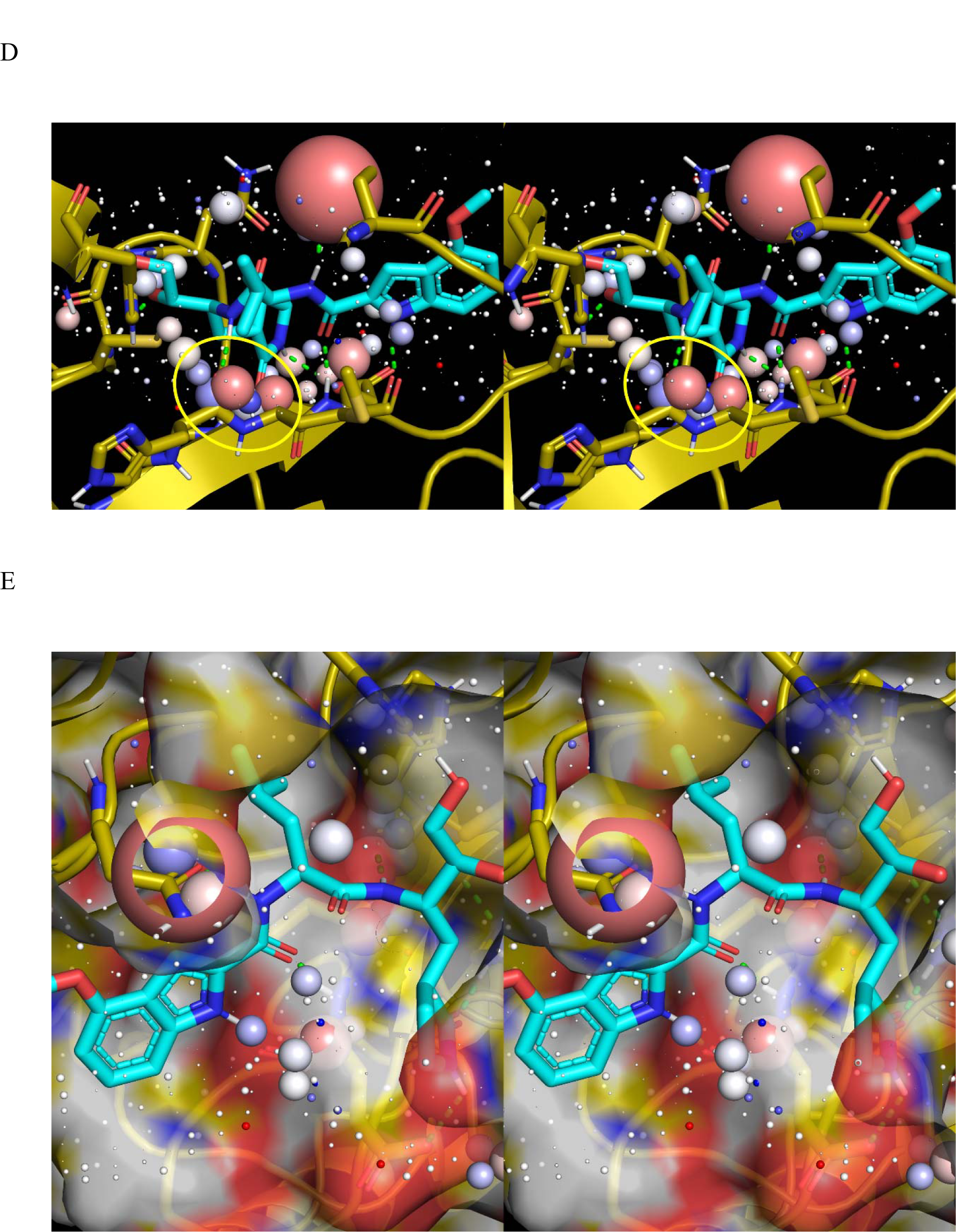
(A) Stereo view of PF-0083521 extracted from 6XHM overlaid on the <u>inverse</u> <u>solvophore</u> of apo monomeric M^pro^ (PDB code = 2QCY), showing protein-inhibitor H-bond replacements for the six high occupancy/H-bond enriched M^pro^ voxel clusters circled in magenta, orange, blue, green, lime green, and yellow (voxel annotations explained in Materials and methods). Protein-inhibitor water H-bond replacements are shown as green dotted lines (noting that the H-bond between His163 and S1 lactam oxygen is hidden by the voxels within the lime green circle). The H-bond between Gln189 and HN2 replaces the high occupancy <u>inhibitor</u> voxels shown in C. We postulate that viral resistance to PF-0083521 is achieved via mutation of Gln189 to amino acids that are <u>incapable</u> of desolvating the HN2 position of the inhibitor (noting that large voxels are fortuitously absent in PF-07321332). (B) Same as A, except showing the solvent accessible surface of PF-0083521. All M^pro^ voxels within the bounds of this surface are desolvated. (C) Front stereo view of apo monomeric M^pro^ overlaid on the solvophore of PF-0083521, showing the protein-inhibitor water H-bond replacements for the four high occupancy voxels circled in red and magenta. The protein-inhibitor H-bonds are shown as green dotted lines. The large voxel at HN2 is replaced by Gln189 (cutaway view). (D) Same as C, except viewed from the rear, showing the protein-inhibitor water H-bond replacements for the chain of high occupancy voxels proximal to HN3 (yellow circle). (E) Same as D, except showing the solvent accessible surface of M^pro^. All of the inhibitor voxels within the bounds of this surface are desolvated.

**Figure 13.**
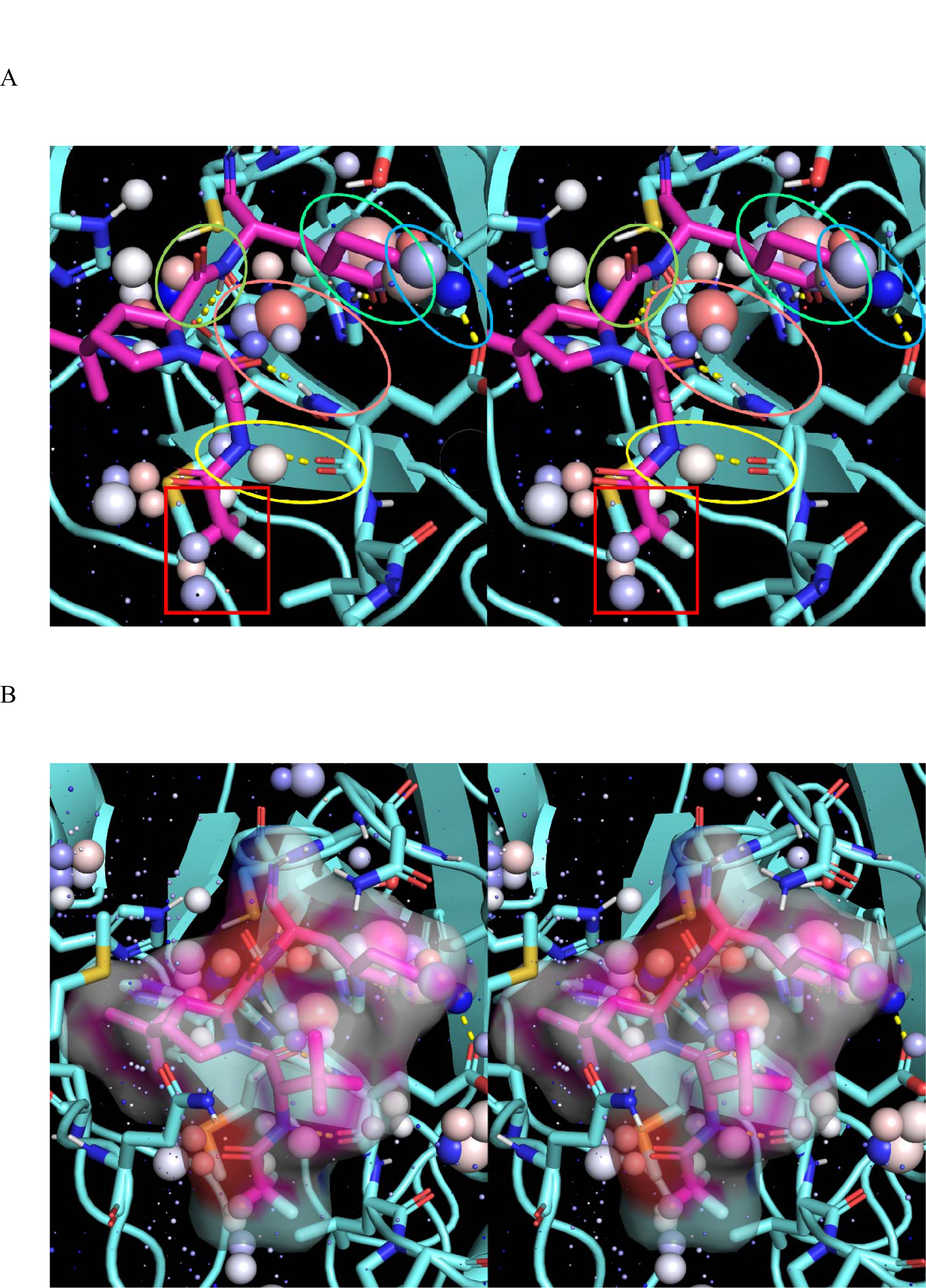
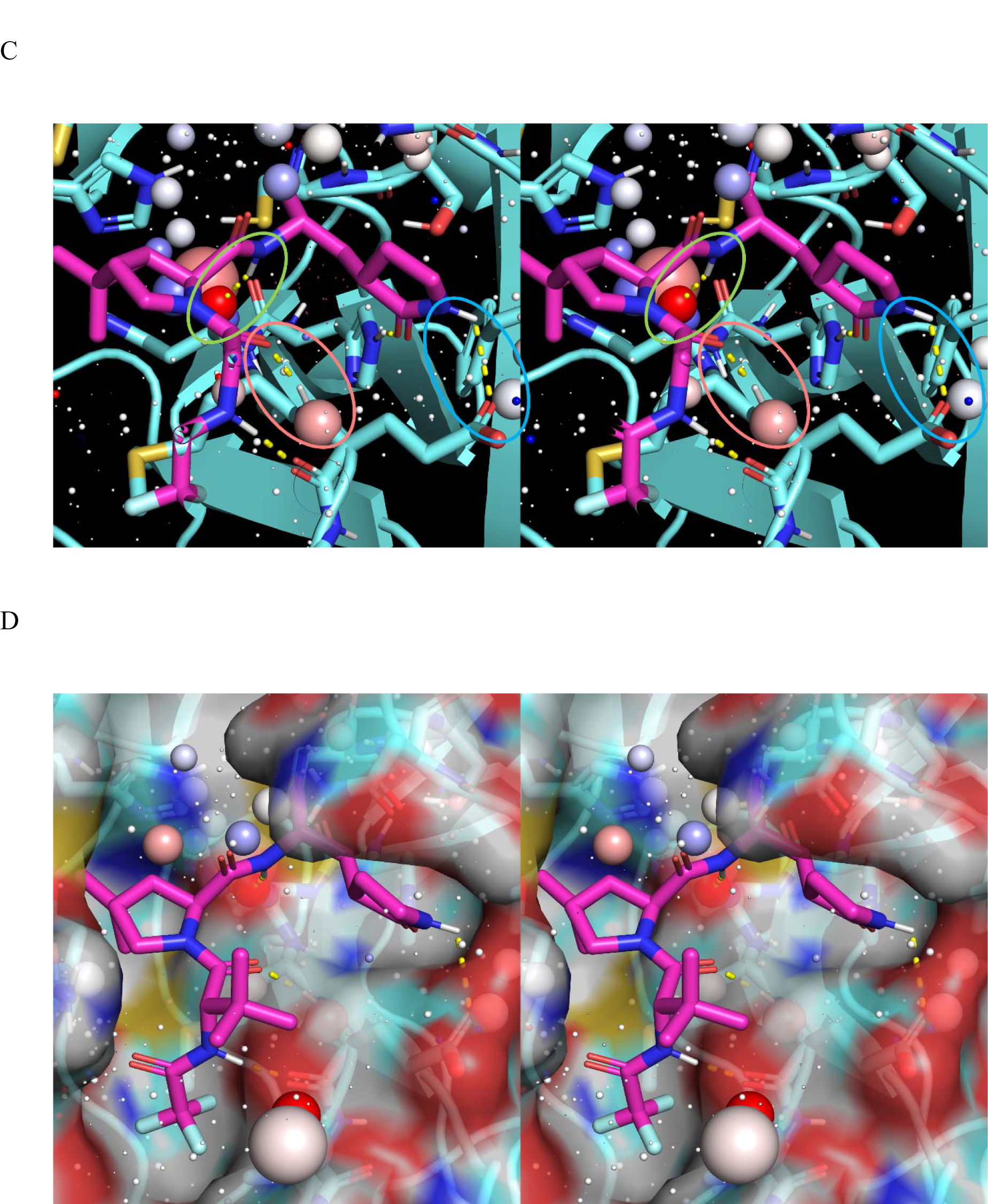
(A) Stereo view of PF-07321332 overlaid on the <u>inverse solvophore</u> of apo monomeric mutant M^pro^, showing the protein-inhibitor H-bond replacements for the five high occupancy/H-bond enriched M^pro^ voxel clusters circled in, orange, blue, green, lime green, and yellow (voxel annotations explained in Materials and methods). Slowed k_on_ is expected due to mismatching between the inhibitor CF_3_ group and the high occupancy voxel cluster enclosed in the red rectangle. The protein-inhibitor H-bond water replacements are shown as yellow dotted lines. (B) Same as A, except showing the solvent accessible surface of PF-07321332. All the M^pro^ voxels within the bounds of this surface are desolvated. (C) Stereo view of apo monomeric M^pro^ overlaid on the solvophore of PF-07321332, showing the protein-inhibitor water H-bond replacements for the three high occupancy voxels circled in orange, green, and blue. (D) Same as C, except showing the solvent accessible surface of M^pro^. All the inhibitor voxels within the bounds of this surface are desolvated.

## Discussion

Improving P_d_ depends on a first principles approach to drug design, in which solubility, permeability, drug-target binding kinetics, and drug-off-target binding kinetics are addressed holistically according to the predicted distributions of H-bond enriched and depleted solvation of hit and lead compounds, target and off-target binding sites, and membrane surfaces (Figure 14) (ultimately aimed at “reverse-engineering” predicted inverse solvophores into LMW compounds capable of achieving optimal PK-PD relationships in humans). Our WATMD results for the COVID M^pro^ inhibitors PF-07321332 and PF-008352 suggest that these compounds were wittingly or unwittingly designed based on these principles. Solubility, as reflected in the distribution and magnitude of H-bond enriched and depleted solvation is arguably more useful for drug design purposes than scalar solubility measurements. Non-covalent binding and membrane permeation are treated in terms of the separate 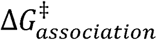 and 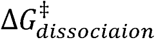 barriers consisting principally of the desolvation and resolvation costs incurred during rearrangements (where higher desolvation costs equate to lower solvation free energy and proportionately higher solubility at the expense of higher maximum) versus the aggregated contributions to the equilibrium ΔG (as reflected in the K_d_, IC_50_, etc.).

**Figure 14.**
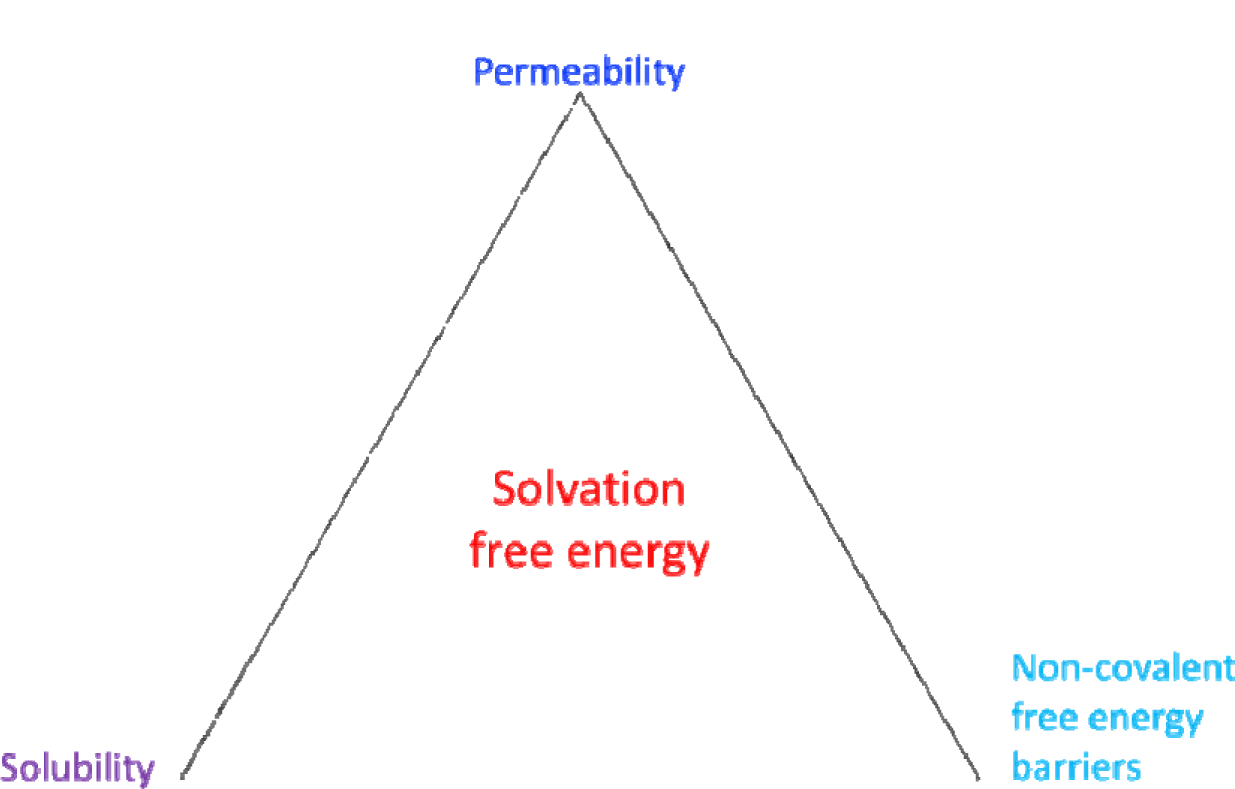
Solubility, permeability, and non-covalent intra- and intermolecular free energy barriers are governed principally by water-solute and water-water H-bond free energy contributions, which are spatially distributed over the solvent accessible surfaces of solutes. Of particular interest are regions of H-bond enriched and depleted solvation versus energetically neutral bulk-like solvation. Solubility (excluding dissolution) is a complex function of the relative proportions, magnitudes, and spatial distributions of H-bond enriched and depleted solvation at polar and non-polar solute surfaces, respectively (enhanced by the former and diminished by the latter). Solvation free energy, in turn, depends on the local H-bonding environment at each solute surface position (governed by the local chemical composition and surface shape/curvature (Wan et al., 2021a)). and are proportional to the total desolvation cost of H-bond <u>enriched</u> solvation during entry or association, respectively, and and are proportional to the total resolvation cost at H-bond <u>depleted</u> positions during exit or dissociation. The maximum or increases with increasing solubility and total desolvation cost (wherein replacement of the H-bonds of H-bond enriched solvation needed to speed k_on_ becomes increasingly challenging), and the maximum 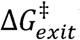 and 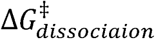 increases with decreasing solubility and total resolvation cost. Permeability is governed by 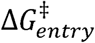 and 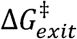 to/from phospholipid membranes, respectively (noting that partitioning occurs when the rate of entry exceeds the rate of exit). The skewed distribution of H-bond enriched and depleted solvation of amphipathic, phospholipid-like solutes promotes membrane partitioning rather than permeation. The Goldilocks zone of H-bond enriched and depleted solvation governing optimal solubility, target binding, and permeability is the underlying basis of the Lipinski Rule of 5 (expressed as polar/non-polar composition) (Lipinski, 2000).

According to our theory, <u>the minimum free energy conformational states of LMW species</u> are those tipped toward excess H-bond enriched versus H-bond depleted solvation (trapped or impeded <u>internal</u> H-bond depleted solvation in particular). The most populated sterically accessible conformational states i of a given species under non-equilibrium conditions are those with the fastest rates of entry (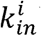), and slowest rates of exit (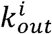) versus those corresponding to free energy minima under equilibrium conditions. The fastest 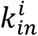 s are those with the lowest <u>desolvation</u> costs (i.e., in which H-bond enriched solvation is replaced by intra-solute H-bonds during entry or in which such solvation is minimal). The slowest 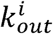 s are those with the highest <u>resolvation</u> costs (i.e., those in which large amounts of H-bond <u>depleted</u> solvation are expelled during entry). The actual conformational distributions of LMW species in solution may be considerably wider than those suggested by force-field-based calculations in which favorable enthalpic interatomic contributions are overestimated. The preferred conformations are likely those in which the overall solvation free energy and electronic energy in the case of unsaturated or covalently rearrangeable moieties (e.g., tautomers) is minimized, as follows:

1. Extended conformations of highly polar compounds in which the polar groups are maximally solvated.
2. Conformations in which the H-bonds of H-bond enriched solvating water are optimally replaced by <u>intramolecular</u> H-bonds (the rates of formation of which depend on the replacement quality vis-à-vis the desolvated H-bond enriched water). Such conformations are transient.
3. Folded conformations in the case of larger solutes, in which H-bond depleted solvation is maximally (though incompletely) desolvated. The persistence of such conformations is proportional to the resolvation cost incurred during exiting.

Desolvation and resolvation costs are treated in terms of their individual (vectorial) contributions at the internal and external surface positions of HMW solutes and external surface positions of LMW solutes, versus conventional measured and calculated <u>scalar</u> quantities under the status quo paradigm (e.g., binding constants, binding free energy, solubility, permeability, logP) (Figure 15).

**Figure 15.**
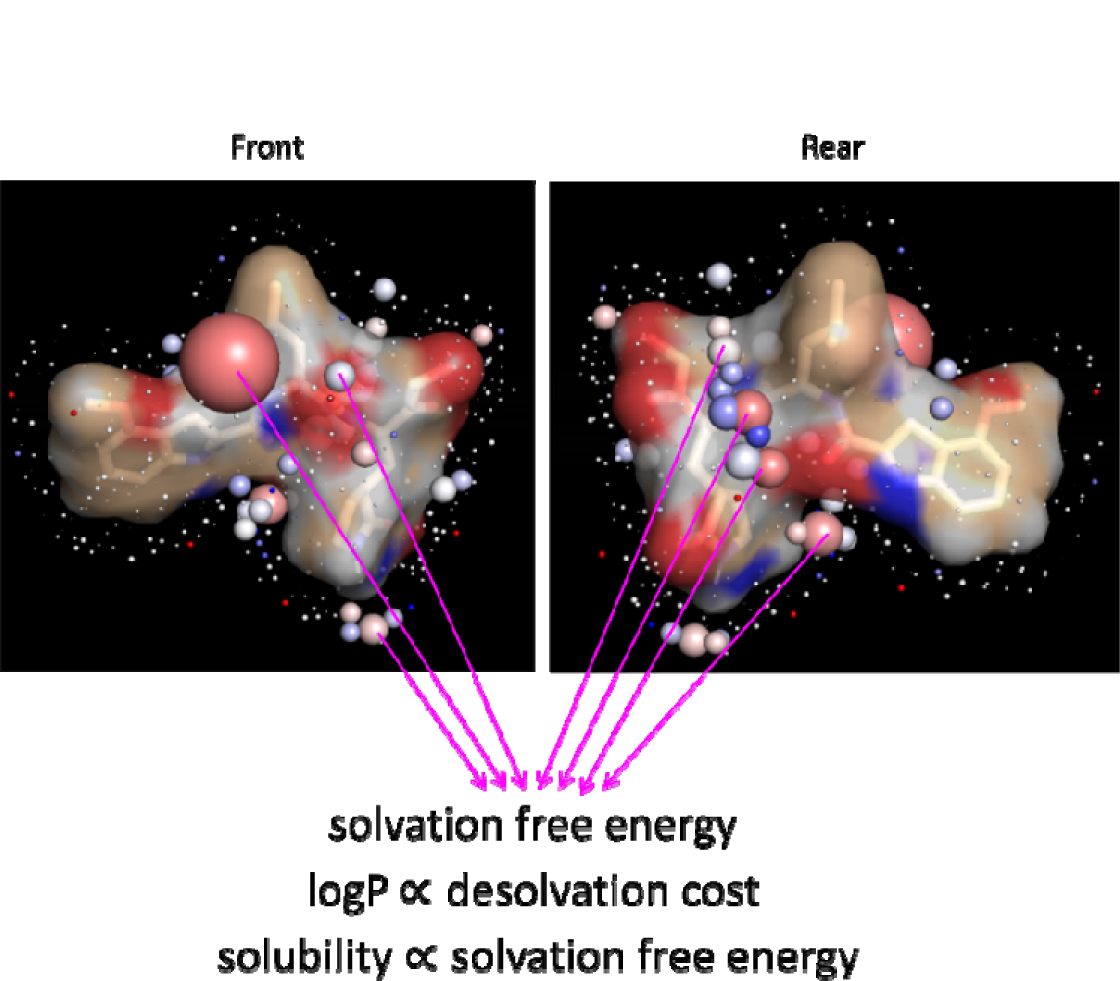
The high occupancy voxels of the M^pro^ inhibitor PF-0083521 calculated using WATMD (see below) overlaid on the solvent accessible surface of the compound (showing the front and rear faces). The total desolvation cost (i.e., -(the total solvation free energy)) consists of the sum of the desolvation costs at each voxel position (reflected in the voxel radii reflecting the time-averaged water occupancy relative to bulk solvent). LogP provides an experimental benchmark of the relationship between the voxel radii and the desolvation free energy (noting that the spatial distribution of the individual contributions to the desolvation and solvation free energies is more useful for drug design than measured scalar logP and solubility values).

Furthermore, target sequence ➔ drug approaches are precluded by the inherently non-linear nature of drug discovery/design resulting from the mutual interdependence of drug properties and behaviors under native cellular conditions (Figure 16).

**Figure 16.**
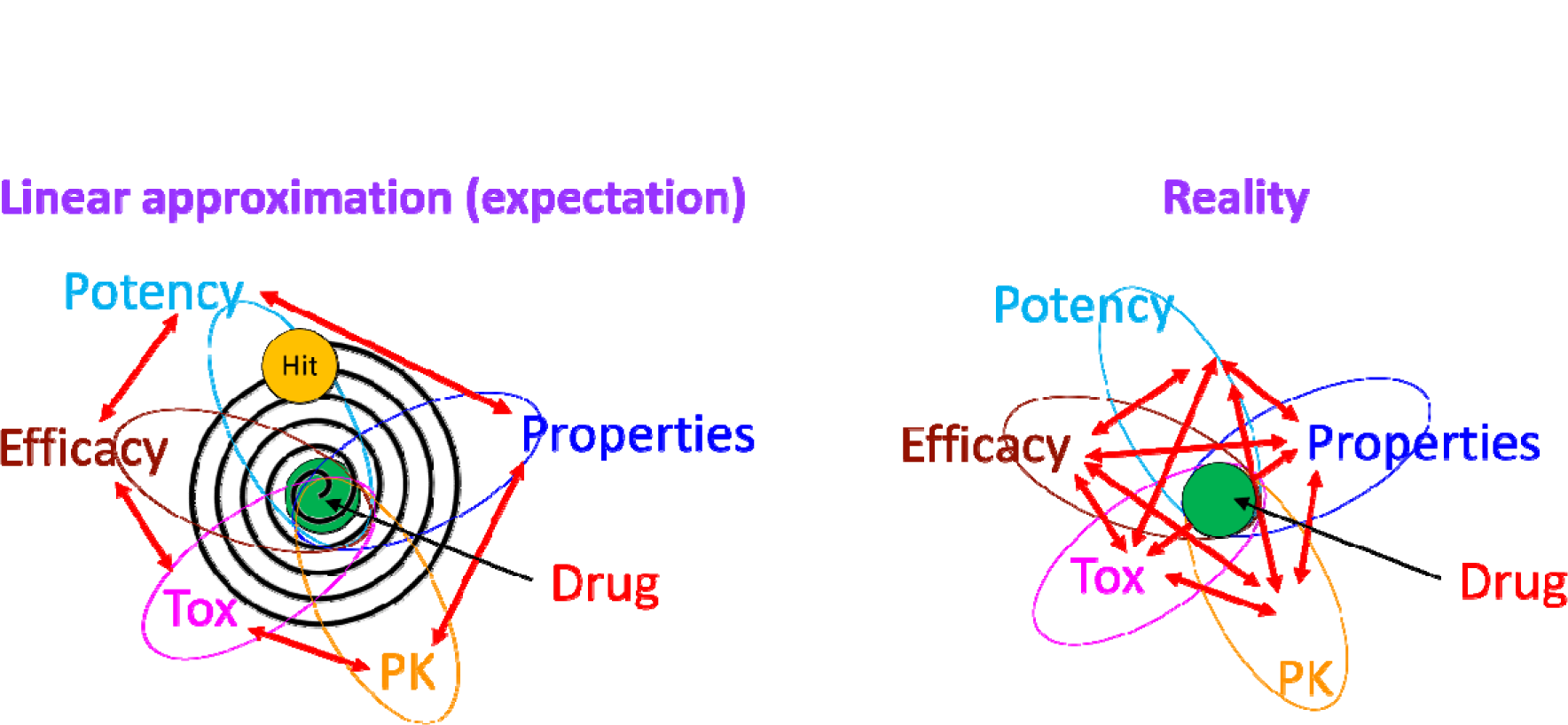
Left: a linear relationship among the major drug behaviors/properties is tacitly assumed under the status quo lead optimization paradigm. Predicted properties are optimized in a step-wise fashion, beginning with chemical starting points (hits) that “spiral” into predicted drugs via cycles of trial-and-error synthesis and testing. Right: in reality, such relationships are highly non-linear due to their mutual interdependence, wherein optimization of one behavior may result in deoptimization of one or more others.

Based on our theory, lead optimization is ideally aimed at:

1. Minimizing the need for dose escalation during clinical testing (which erodes the safety margin and TI) by tuning k_on_ and k_off_ to the rates of target (or target binding site) buildup and decay, respectively (Selvaggio and Pearlstein, 2018) such that the minimum γ_eff_ in humans is achieved at the lowest possible free C_max_. Covalent reactions can be used to slow k_off_ when the amount of H-bond depleted solvation is insufficient for achieving the minimum efficacious γ_eff_ (noting that the occupancy of such drugs accumulates to the equilibrium level over time).
2. Achieving the Goldilocks zone of solubility, permeability, and drug-target and drug-off-target binding, in which the overall desolvation costs are sufficiently low (reflected in higher logP). The optimal solvation properties correspond to:

a. The Goldilocks zone of solubility and desolvation costs incurred during drug-target and drug-membrane association (Figure 17). Poorly soluble compounds dissolved in DMSO stock solutions (frequently used in assays) revert to their aqueous solubilization level and may distribute among solubilized (the bioavailable fraction), suspended, and precipitated fractions.

**Figure 17.**
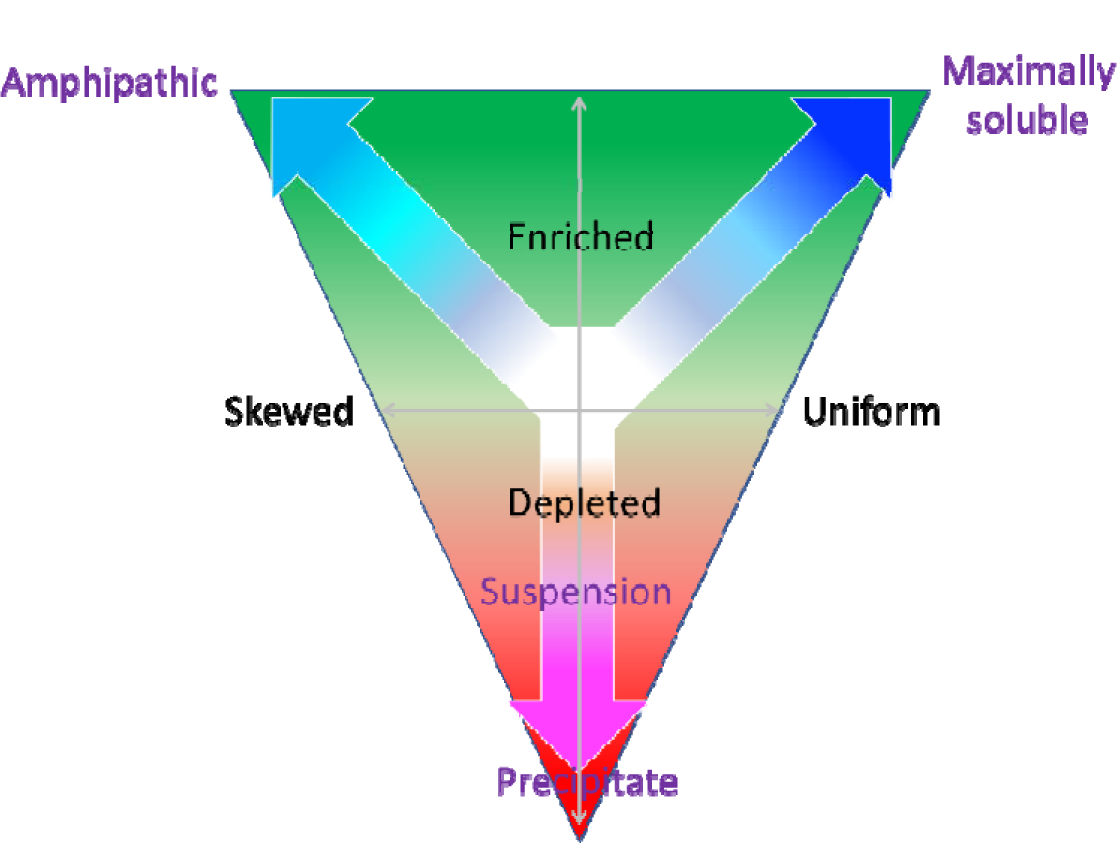
Maximal solubility depends on uniformly distributed H-bond enriched solvation across solute surfaces, which scales downward as the ratio of H-bond enriched/depleted solvation and the absolute free energy magnitudes thereof decrease. The asymmetric distributions of H-bond enriched and depleted solvation in amphipathic molecules are exaggerated by acid groups and basic amines. Precipitation occurs at a high threshold ratio of H-bond depleted to enriched solvation, whereas suspensions occur at a lower threshold ratio.
b. Sufficient <u>polar groups</u> optimally positioned for low-cost desolvation of the H-bond enriched water solvating the polar phospholipid groups at membrane surfaces and target binding sites (avoiding mismatches between non-polar drug groups and H-bond enriched solvation), resulting in fast k_on_ and membrane k_in_. Fast k_on_ is predicted for PF-0083521 and PF-07321332 on this basis (see the results section).
c. Sufficient <u>non-polar groups</u> optimally positioned for desolvating H-bond depleted solvation within the target binding site (avoiding mismatches between polar drug groups and H-bond depleted solvation), resulting in slow k_off_ (noting that excess H-bond depleted solvation results in poor solubility, and in some cases, membrane partitioning (i.e., slow membrane k_out_). Slow k_off_ is predicted for PF-0083521 and PF-07321332 on this basis (see the results section).
d. Gains in H-bond <u>enriched</u> solvation <u>in the bound state</u> (which slows k_off_) but not H-bond <u>depleted</u> or trapped/impeded solvation (which slows k_on_).
e. Avoiding basic groups that are frequently used to solubilize poorly soluble compounds. The solvation of such compounds is distributed asymmetrically between H-bond enriched solvation of the basic group and H-bond depleted solvation of the remaining scaffold (translating to amphipathicity/quasi-amphipathicity). Excess H-bond depleted solvation relative to enriched (the direct cause of poor solubility) can result in non-specific binding, membrane partitioning (i.e., a high volume of distribution). Basic amines can further result in high levels of lysosomal trapping (Llanos et al., 2019) and ion channel blockade.
f. The lowest possible molecular weight distributed between polar and non-polar groups required for matching H-bond enriched solvation on membrane and target binding site surfaces and depleted solvation on target binding site surfaces, respectively (the Goldilocks zone of chemical composition). This principle is captured in qualitative scalar form by the Lipinski rule of 5 (Lipinski, 2000).

Activity cliffs (abrupt changes in potency due to small chemical changes) are due to significant k_on_ slowing, significant k_off_ speeding, or both. Slowed k_on_ relative to the parent is symptomatic of the loss of H-bond free energy due to:

1. Suboptimal H-bond replacement of H-bond <u>enriched</u> solvation of the modified analog and/or binding site relative to the reference analog, stemming from altered geometry and/or chemical composition of the offending R-group (translating to increased desolvation cost).
2. The generation of additional H-bond <u>depleted</u> solvation in the bound state of the modified analog compared with the parent, which destabilizes the bound state of the analog relative to that of the parent.

Sped k_off_ is symptomatic of the loss of H-bond free energy among the partners (i.e., decreased resolvation cost) due to:

1. Reduced desolvation of H-bond <u>depleted</u> solvation of the modified analog and/or the binding site caused by spatial mismatching between the offending R-group and binding site relative to the parent.
2. The loss of additional H-bond <u>enriched</u> solvation generated in the bound state of the parent but not the analog, resulting in destabilization of the bound state of the analog relative to that of the parent.

## Conclusion

Failure to achieve a TI during clinical testing in appropriate patient populations stems from poorly predicted therapeutic effects of drug-target occupancy, structure-free energy relationships governing exposure-target occupancy (PK-PD) relationships, and/or overlapping minimum efficacious and toxic exposure levels in humans that may be attributed in some cases to kinetically mistuned drug-target binding. Structure-activity relationship models obtained via conventional statistical methods or deep learning are generally useful for capturing weak signals in noisy datasets (e.g., protein sequence-tertiary/quaternary structure relationships in Alphafold2 (Jumper et al., 2021)), but are subject to disconnects between the dependent and independent variables (e.g., measured drug-target binding data and molecular descriptors based on interatomic contacts) and between disconnected contexts (e.g., equilibrium conditions in vitro versus non-equilibrium conditions in the native cellular setting). Predictions based on first principles theoretical understanding (the rules of the game) are far more likely to improve preclinical and clinical outcomes than empirical data models. Biodynamics-guided drug design is concerned with achieving the Goldilocks zone of solvation free energy contributions to solubility, permeability, and dynamic drug-target fractional occupancy (i.e., not too H-bond depleted and not too H-bond enriched). Improving preclinical and clinical success rates depends to a large degree on shifting the paradigm from trial-and-error, process-driven technology/chemistry/screening-centric workflows revolving around equilibrium data and interatomic contact models to one guided by scientific principles that are better aligned with the native non-equilibrium cellular setting.

